# In silico optimization of deep brain stimulation to enhance cognitive control: Improving performance and practicality with a continuous rolling arena

**DOI:** 10.64898/2026.08.20.745844

**Authors:** Sumedh S Nagrale, Alik S Widge

## Abstract

**Objective:** The use of Deep Brain Stimulation (DBS) on the ventral capsule/ventral striatum (VCVS) has therapeutic potential for patients with refractory psychiatric disorders, but clinical success is impeded by the need for a time-consuming and trial-and-error process when setting the parameters, this process relying on subjective self-reports. By monitoring objective behavioural markers it is possible to quickly assess the contact settings. In this study, we assess a direct closed-loop Multi-Armed Bandit (MAB) optimisation framework which is designed to quickly determine the best stimulation contacts by using raw reaction times (RT) obtained during a cognitive control task.

**Approach:** We leveraged empirical data demonstrating that VCVS DBS enhances cognitive control during the Multi-Source Interference Task (MSIT) in a site-specific manner. Using a synthetic patient simulation environment across 1,000 replicates, we benchmarked adaptive MAB algorithms under noisy, non-stationary conditions. Crucially, we eliminated intermediate state-space sensor models to evaluate raw RT directly, transitioned from discrete daily resets to an uninterrupted continuous optimization architecture, and implemented a rolling arena mechanism to scale contact selection under real-world hardware constraints.

**Main results:** Eliminating the intermediate sensor model prevented high-frequency noise amplification (where state variance was inflated by 51.7% in baseline and 148.0% in conflict states) and reduced contact ranking failure rates from 31.9% down to 11.9%. Operating within a continuous trial architecture preserved historical sample density, driving mean trial-level regret down steadily over 4,200 trials and enabling dynamic re-convergence across unannounced mid-session change-points. Additionally, a 4-contact sub-arena successfully scaled search efficiency across 8-contact arrays without sacrificing selection accuracy.

**Significance:** Direct MAB optimization within a continuous rolling arena provides a noise-resilient, hardware-compatible architecture for automated DBS programming. By bypassing latent state estimation and utilizing standard task-based behavioral metrics without specialized recording hardware or complex state-space modeling, this framework reduces search timelines to clinically feasible durations, establishing a scalable foundation for real-time, patient-tailored neuromodulation.

## I. Introduction

Psychiatric disorders constitute a major global health burden, contributing to long-term disability, economic loss, and diminished quality of life. Standard therapeutic regimens primarily rely on pharmacotherapy such as selective serotonin reuptake inhibitors (SSRIs) combined with psychotherapy modalities like cognitive behavioral therapy (CBT). However, approximately one-third of patients exhibit treatment-resistant psychiatric conditions, failing to achieve sustained symptomatic relief. For this refractory subpopulation, advanced neuromodulation technologies have emerged as effective alternative interventions.

Deep Brain Stimulation (DBS) for psychiatric disorders is a surgical neuromodulation therapy involving the stereotactic implantation of leads into specific subcortical structures. These leads deliver continuous, 130 Hz high-frequency electrical pulses to disrupt pathological neural activity and modulate dysfunctional cortico-striatal circuits. While non-invasive approaches like transcranial magnetic stimulation (TMS) offer broader cortical stimulation, DBS provides finer spatial precision and deep subcortical access. Consequently, DBS has been actively investigated across several key anatomical targets for treatment-resistant depression (TRD) and obsessive-compulsive disorder (OCD) to engage networks implicated in dysfunctional cognitive and affective processing. These targets include the ventral capsule/ventral striatum (VC/VS), the subgenual cingulate cortex (SCC), and the superolateral medial forebrain bundle (slMFB). Among these targets, the VC/VS has demonstrated key clinical significance, receiving US Food and Drug Administration (FDA) Humanitarian Device Exemption (HDE) approval for OCD and serving as the only psychiatric DBS target to successfully meet its primary endpoint in a randomized, sham-controlled trial for MDD[1].

Despite its established therapeutic potential, DBS across these clinical targets remains limited by modest short-term response rates. In double-blind, sham-controlled VC/VS DBS studies, response rates typically range from 20% to 40% [2–6], although these rates are significantly higher, closer to 70% in open-label data [3,7]. Similar response rate limitations are observed across other clinical targets, such as the SCC [8] and the slMFB [9,10]. While response rates often improve to 50-60% during long-term, open-label studies these gains are achieved through a tedious parameter titration process [1,11].

Along with post-surgical anatomical variability across participants, a primary driver of therapeutic delay is the selection of stimulation parameters, i.e. DBS programming. Currently, this is a trial-and-error process of identifying optimal settings within a vast search space. These parameters include contact selection, amplitude, frequency, and pulse width. For instance, [1] required a mean 51.6 week optimization phase involving biweekly adjustments across four contacts along the ALIC gradient. This extended evaluation period persists because parameter tuning relies on subjective, episodic scales (e.g., standardized rating scales like HAM-D-17 or MADRS, alongside qualitative, acute mood reports) rather than real time physiological feedback. Further, evaluating each parameter configuration takes weeks. These subjective scales are inherently noisy, time-delayed and subject to high placebo response or fluctuating variance. Furthermore, these subjective stimulation responses can depend heavily on externally influenced states [12–14].

To improve parameter optimization, recent approaches have leveraged advanced imaging. These methods use connectomic electric-field modeling to map white matter bundle engagement based on electrode placement [4,15,16]. Although the structural imaging confirms anatomical target it can not determine whether the connected network responds functionally, and as such, tract modeling has been inconsistent in predicting clinical response [17]. Alternatively, direct neurophysiological sensing such as local field potentials (LFPs) or cortico-cortical evoked potentials (CCEPs) has been proposed to monitor real time network engagement [12,18,19]. Although this approach is successful in movement disorders, identifying reliable physiological markers for DBS tuning in non-motor applications has proven difficult again owing to the poor signal-to-noise ratio of self-report and a lack of quantitative, predictive behavioral assays and reliable neural biomarkers [18–20].

To bridge this gap, a more scalable approach is to evaluate circuit engagement indirectly through behavior. Modulating these targeted circuits produces rapid, measurable changes in specific brain functions [21]. For example, DBS for tremor is titrated in the clinic by monitoring motor suppression in real time. Similarly, psychiatric stimulation parameters can be continuously optimized by tracking objective behavioral metrics, such as performance on standardized psychophysical tasks.

A key behavioral metric for psychiatric circuits involving the ALIC and connected prefrontal cortex (PFC) regions is cognitive control. Cognitive control (CC) is the ability to suppress a prepotent response in favor of a goal-oriented action. It represents a key target for behavioral optimization because cognitive control deficits are common across many psychiatric disorders [22,23]. As a proof of concept, human studies have demonstrated that high-frequency (~130 Hz) VC/VS stimulation rapidly enhances cognitive control, quantitatively measured by Reaction Time (RT) performance during standardized cognitive conflict tasks such as the Multi-Source Interference Task (MSIT) [20,24]. Unlike traditional clinical rating scales that require weeks or months to reflect therapeutic changes, the behavioral effects of stimulation begin and end within seconds of parameter changes on the MSIT. Furthermore, this effect is highly site-specific. Cognitive control improvement varies across individual electrode contacts. Even millimeter-scale shifts in the stimulation location produce distinctly different behavioral outcomes [24].

Recent reverse-translational research in rodents confirmed these human findings using mid-striatal DBS [25]. Specifically, computational modeling revealed that stimulation selectively accelerates the rate of evidence accumulation without inducing motor hyperactivity, impulsivity, or accuracy tradeoffs. Furthermore, modeling confirmed that this same mechanism operates in humans. Consequently, task-based cognitive control offers a rapid, objective, and cross-species biomarker to guide DBS parameter titration. At the circuit level, preclinical optogenetics demonstrated that activating prefrontal-originating axons, rather than local striatal cells, drives this cognitive control improvement [26]. High-resolution tracing identified the responsible white-matter fibers, which originate specifically in the dorsolateral prefrontal and anterior cingulate cortices. Translating these anatomical insights to humans, patient-specific diffusion tractography links these specific pathways to rapid behavioral changes, providing a clear framework to explain individual differences in stimulation response [27]. Together, these findings establish task-based cognitive control as a scalable, mechanistically grounded biomarker for closed-loop parameter optimization.

However, stimulation-induced RT changes are subtle (typically 5% to 10% of the overall scale) and masked by trial-to-trial behavioral variability. As a result, human clinicians cannot manually detect these signals or distinguish between adjacent contacts in real time. To overcome this limitation, recent work introduced adaptive computational frameworks to automate parameter optimization. Specifically, [28] formulated contact selection as a discrete Multi-Armed Bandit (MAB) decision problem, treating each electrode contact as an individual “arm” with an unknown behavioral reward distribution. By pairing this framework with an Upper Confidence Bound (UCB1) algorithm, this approach systematically resolves the fundamental exploration-exploitation trade-off. It strategically samples uncertain contacts to quantify their effect (exploration) while repeatedly testing high-performing contacts to confirm genuine circuit engagement (exploitation). In silico validations demonstrate that a UCB1-driven MAB framework successfully filters stochastic behavioral noise, achieving reliable convergence to an optimal contact setting within a clinically feasible testing window [28].

While our previous closed-loop system established the feasibility of using a multi-armed bandit algorithm (UCB) and RT-based feedback to identify optimal stimulation contacts, clinical translation requires a more rigorous characterization of algorithm performance. Specifically, algorithms must be evaluated under challenging and realistic operating conditions, assessed using metrics that capture both accuracy and optimization efficiency (e.g., regret), and analyzed to identify the factors that drive successful or unsuccessful convergence. Building on the proof-of-concept findings, this study focuses on the key developments required to move toward a clinically viable optimization framework.

First, we establish that direct reaction-time (RT) feedback outperforms latent state-space modeling for closed-loop optimization. The sensor model was originally designed to achieve two goals. First, it estimates underlying cognitive states from reaction times (RT). Second, it filters out sudden behavioral noise. We estimated this model using the Expectation-Maximization (EM) algorithm. However, our analysis reveals a major practical limitation. While the EM-based model tracks overall cognitive changes very accurately, with a normalized root-mean-square error (NRMSE) under 5 to 9%, the decomposition and smoothing effect removes key information and reduces the discriminative information available to the optimization algorithm, resulting in lower decision accuracy than approaches based directly on RT measurements. These findings demonstrate that, despite accurate signal tracking, the sensor model estimation artifacts degrade decision accuracy and suggest that future use of latent cognitive-state representations will require more advanced estimation approaches.

Second, to ensure a fair comparison across algorithms, we optimized and tuned each algorithm to operate at its best. To achieve this, we tuned the hyperparameters using a comprehensive grid search, and we evaluated different prior distributions to identify the structure best suited for the noisy RT biomarker. This calibration ensures that our evaluation reflects the true upper bounds of each algorithmic approach.

Third, we tested a continuous trial architecture rather than splitting the simulation into discrete, day-wise blocks of 600 trials. This continuation protocol preserves sequential sample histories, ensuring all algorithms maintain identical historical data density. This approach prevents the destructive reset of past performance data, which directly benefits algorithms that learn dynamically from cumulative data. Crucially, a larger pool of sequential trials allows the algorithms to identify noise more effectively. As the total trial count increases, the estimation of the reaction time (RT) improves over trials. This variance reduction provides the optimization framework with a higher-fidelity signal, ultimately enhancing the overall decision-making performance of the system.

Fourth, we expanded the performance metrics to better understand algorithmic behavior under varying constraints. Prior work’s decision accuracy only measures the final outcome. To gain a better understanding, we evaluate both trial-level and decision-level regret across different conditions. While accuracy shows whether an algorithm eventually finds the correct settings, regret metrics reveal the path taken to get there. Specifically, regret analysis demonstrates which frameworks minimize suboptimal choices early on, which ones consistently lock onto the true optimal point, and which ones are prone to random exploration. This allows us to map specific algorithmic strengths to distinct clinical and system priorities.

Fifth, we address the critical practical constraint of a limited trial budget. In standard Multi-Armed Bandit (MAB) problems, algorithms assume a high trial volume to guarantee convergence. However, human patients can only complete a limited number of behavioral trials before fatigue sets in and this is also limited by clinical hours available. To investigate this limitation, we evaluate four distinct convergence criteria: arm selection stability, posterior variability, regret flattening, and a fixed trial threshold. The effectiveness of each criterion depends heavily on how a given algorithm searches the parameter space. For exploitation-heavy frameworks, these criteria allow for early termination, thereby conserving valuable patient trials. Conversely, highly exploratory algorithms require more extended sampling windows to achieve stability. Ultimately, this analysis helps determine how to minimize patient burden without sacrificing optimization accuracy.

Sixth, we execute a rigorous stress test to evaluate algorithmic resilience across a spectrum of signal-to-noise ratios (SNRs). Prior work evaluated these noise levels strictly within a day-wise optimization framework. However, because our new continuous trial architecture provides a much larger pool of sequential data, algorithms that rely on cumulative learning should theoretically perform better over longer runs. Since the precise clinical noise level in real-world patients is unknown, we evaluate performance against simulated ground truth data across varying noise levels. Additionally, to understand the robustness and ability of these algorithms to track sudden changes in the stimulation effect on RT, we perform a change-point analysis where we abruptly alter the underlying effect parameters. Although this represents a highly theoretical scenario, it provides a critical marker of algorithmic robustness. Identifying a framework that maintains high decision accuracy, exhibits low regret across all noise conditions, and dynamically tracks underlying cognitive changes reveals the most practical solution for clinical translation.

Seventh, we address a major hardware constraint in clinical translation: the limited number of stimulation sites that can be optimized simultaneously. While simulations often evaluate fixed problem sizes (e.g., 2, 4, 6, or 8 contacts), real-world deployments are restricted by hardware bandwidth and safe patient trial budgets. To overcome this limitation, we propose a dynamic rolling arena tournament method. The system configures the active optimization space to four stimulation contacts at a time; it systematically eliminates the worst-performing option and rolls in a new, untested electrode contact. This iterative approach ensures the framework remains highly sample-efficient and practically extensible to an arbitrary number (n) of electrode contacts without sacrificing selection accuracy.

The remainder of this paper is organized as follows. Section II details our methods, starting with the updated closed-loop system architecture without the sensor model, followed by the mathematical formulation of the patient generator model used for simulations. It then outlines the core multi-armed bandit optimization algorithms, and defines the trial-level and decision-level performance metrics alongside our proposed convergence criteria and practical scaling rolling arena tournament architecture. Section III presents the empirical results, systematically evaluating the elimination of the sensor model, continuous optimization dynamics, algorithmic calibration, and environmental stress-testing across various noise thresholds. Section IV discusses the clinical-engineering trade-offs of volatility, outlines safety implications for real-world translation, and addresses the limitations of the study. Finally, Section V concludes the paper by framing these developments as translational outcomes for closed-loop neurostimulation.

## II. Methods

### Closed Loop System Architecture

The closed-loop optimization framework consists of two main elements: a Generator, which combines a generative patient environment with a simulated behavioral task, and a decision-making multi-armed bandit algorithm (Figure 1). This core architecture builds upon the layout established in our previous work, with a critical modification: the elimination of the latent sensor model. By bypassing this sensor state-estimation layer, the optimization engine transitions from optimizing an estimated cognitive state to operating directly on a behavioral biomarker (RT). At each trial, the bandit algorithm selects a stimulation contact configuration; the Generator then produces a corresponding RT for the task, providing immediate feedback to the algorithm. This iterative cycle continues until a specified convergence criterion is met.

**Figure 1.**
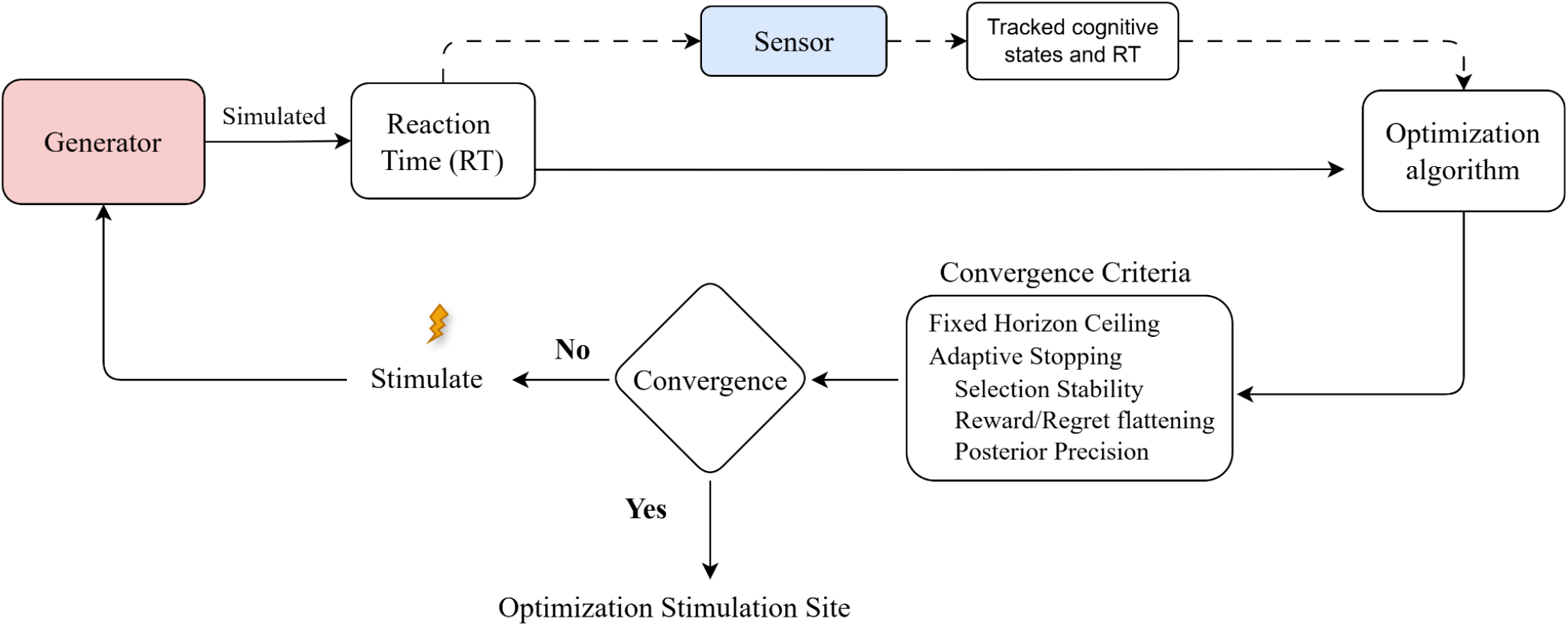
Operational schematic of the closed-loop stimulation optimization architecture. The Generator models patient deep brain stimulation (DBS) response dynamics during the Multi-Source Interference Task (MSIT). Arrows indicate the trial-by-trial flow of information. The Generator internally integrates an MSIT task paradigm (congruent vs. incongruent trials), an autoregressive state-space generative process, and an underlying contact-response surface to output simulated reaction times (RT). In the proposed streamlined architecture, the optimization algorithm receives raw reaction time measurements directly as real-time behavioral feedback (solid arrow). This stands in contrast to prior designs (dashed block/arrow), which routed feedback through an intermediate state-space sensor model to estimate latent cognitive states and smooth RT. Before suggesting a new stimulation contact configuration for the Generator, optimization progress is checked against a Fixed Horizon Ceiling (evaluated across both architectures) and adaptive stopping criteria (evaluated specifically for the direct raw RT design) to determine whether termination or convergence has occurred. Over successive trials, this continuous feedback loop drives the system toward the optimal electrode target.

### Mathematical Formulation of Patient Generator Model

#### 1. The Multi-Source Interference Task (MSIT)

To evaluate the closed loop stimulation optimization paradigm we use the Generator model that replicates the patient’s trial by trial behavior during the Multi-Source Interference Task (MSIT). MSIT actively engages cognitive control networks by presenting spatial and structural interference. MSIT consists of two trial types: low conflict (congruent) trials and high conflict (incongruent) trials. In congruent trials, the target digit aligns directly with its physical response key. In incongruent trials, the target is flanked by distractor digits and spatially mismatched, requiring the network to resolve cognitive interference. Successful performance requires both reactive and proactive cognitive control, measured explicitly through the participant’s reaction time (RT).

#### 2. Empirical Dataset

To develop the simulated patient model, we utilize an empirical dataset of six intracranial monitoring participants (S1–S6) with longstanding pharmaco-resistant epilepsy [24,28].These individuals were originally reported as participants 8-13 in [24]. Participants voluntarily enrolled with fully informed consent obtained by a study staff member who was not the participant’s primary clinician in accordance with guidelines and procedures approved by the local institutional review boards at Partners Healthcare (Massachusetts General Hospital), with secondary review from the US Army Human Research Protections Office.

A 600 ms symmetric biphasic pulse train (130 Hz, 2-4 mA, 90μs pulse width) was delivered at image onset to influence decision-making dynamics. This stimulation was applied via bipolar contact pairs across up to four locations in the ventral and dorsal internal capsule (IC). The total dataset comprises 2,207 MSIT trials across the cohort, with individual trial totals ranging from 320 to 440 per subject (S1: 343; S2: 378; S3: 320; S4: 383; S5: 383; S6: 440). Active stimulation was administered in 1,407 total trials (192–255 trials per participant) alongside 110, 124, 128, 128, 159, and 191 non-stimulation control trials.

#### 3. Data model

During the MSIT, trial-level reaction times (RT) exhibit right-skewed distributions that are accurately modeled using Gamma distributions, whose underlying parameters shift according to cognitive demand (e.g., congruent vs. incongruent conditions). The observed reaction time denoted as (*Z*_*RT*_) is given by

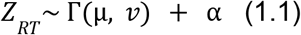

where α > 0 is the offset term representing the minimum bound of the reaction time, 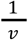 accounts for trial-to-trial response variability, and μ represents the expected mean reaction time, which experiences both task-driven systematic shifts and stochastic fluctuations across trials..

Grounded in prior state-space modeling paradigms [24,28], the mean reaction time μ is decomposed into two separate latent cognitive states connected via a log-link function

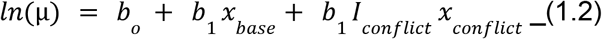

Where *x*_*base*_ reflects core cognitive processing speed in the absence of conflict, and an interference cognitive state (*x*_*conflict*_), measures the supplementary cognitive effort required during high-interference conditions, and *I*_*conflict*_ ∈ {0, 1} acts as a trial-specific conflict indicator flag (0 for congruent, 1 for incongruent). The regression parameters *b*_*o*_, *b*_1_, and *b*_2_ set the structural baseline performance and weight the behavioral contributions of each cognitive state, respectively. The sequential evolution of these states from trial k to k+1 follows coupled first-order autoregressive [AR(1)] processes.

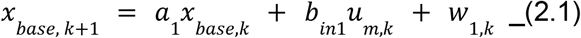

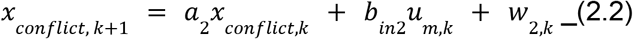

where the decay constants *a*_1_ and *a*_2_ govern state evolution over trials, and 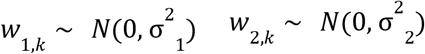 are mutually independent, zero-mean Gaussian state noise processes. To mirror our human clinical environment [24], the binary input vector *u*_*m,k*_ indicates active monopolar deep brain stimulation (DBS) across candidate electrode contact sites. While the gain vectors *b* _*in*1_ and *b*_*in*2_ control how stimulation drives the baseline and conflict states, respectively. Under this state-space formulation, active electrical stimulation does not cause an immediate step-change in observable reaction time, but rather induces a gradual, state-dependent trajectory toward a new steady-state performance level, matching observed human clinical dynamics [24].

#### 4. Generator Model

The generator model reproduces trial-by-trial patient performance during the MSIT by leveraging the state-space formulation in Equations (1.1)–(2.2). Parameter identification and cognitive state estimation are conducted using the COMPASS toolkit, which applies an Expectation-Maximization (EM) algorithm to jointly infer model parameters and latent cognitive trajectories from observable reaction times (RT). In its unconstrained representation, the full parameter vector resides in a high-dimensional search space, which presents a significant risk of convergence to sub-optimal local maxima

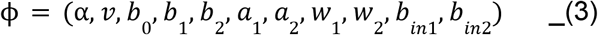

To overcome this dimensionality challenge and ensure numerical stability, structured constraints are imposed prior to model realization. The input gain vectors *b*_*in*1_, *b*_*in*2_ are initially set to zero and subsequently assigned as in “Optimization Problem Simulation and Response Surfaces”. The cognitive state scaling coefficients are fixed to unity (*b*_1_, *b*_2_ = 1), and the initial latent state conditions *x*_0_ in Equations (2.1)–(2.2) are sampled from a standard normal distribution *N*(0, 1). Following validated parameter bounds from prior work [24,28], the autoregressive state persistence coefficients are fixed near unity (*a*_1_ = 0.9999, *a*_2_ = 0.9999) to capture long-term cognitive continuity across trials. To embed physiological behavioral variability across the simulated cohort, the process noise variances *w*_1_ and *w*_2_ are drawn randomly from empirical ranges [0.0039, 0.0884] and [0.00008,0.0110] for *x*_*base*_ and *x*_*conflict*_, respectively. Together, these structural constraints reduce the parameter estimation space to a baseline triplet:

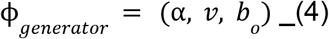

The reduced parameter set in Equation (4) is estimated via Maximum Likelihood (ML)using non-stimulation control blocks from the empirical cohort, with estimation convergence achieved when Δ*ML* < 0. 001. Using this constrained estimation pipeline, a pool of 1,000 synthetic patient profiles (the “generator pool”) was generated for each of the six clinical participants (S1–S6).

#### 5. Sensor Model

In a clinical setting, latent cognitive states cannot be directly observed and must be dynamically reconstructed by fitting the state-space framework to empirical, trial-by-trial reaction times. This state decoding is performed by the sensor model, whose reduced parameter estimation vector is defined by:

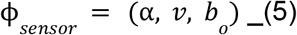

Parameter identification for the sensor model is executed using the COMPASS Expectation-Maximization algorithm with a convergence threshold of Δ*ML* < 0. 001. The process noise variances *w*_1_ and *w*_2_ are initially drawn from empirical bounds [0.0156, 0.0884] and [0.00008,0.0110], respectively, with an additional 0.0625 added to both. This inflated process noise covariance compensates for the omission of an explicit stimulation input term in the filter design, enabling the state estimator to adapt rapidly when active deep brain stimulation (DBS) drives state trajectories away from un-stimulated AR(1) baseline dynamics. For each individual model realization, initial latent state values *x*_0_ in Equations (2.1)–(2.2) are sampled from a standard normal distribution *N*(0, 1). To accommodate broad behavioral heterogeneity across synthetic generator outputs, the reaction time lower bound α is set to 0.1. Using this state-decoding architecture applied to synthetic behavioral outputs from the generator pool, 1000 distinct sensor models were fitted per participant cohort.

While the core experimental analyses throughout this work rely on the architecture without a sensor model, we incorporate the sensor model specifically to evaluate and contrast performance metrics under state-observation constraints. Explicitly testing the system both with and without this state-decoding layer allows us to isolate the impact of unobservable latent dynamics versus direct state access, ensuring a clear baseline comparison before proceeding with the primary experimental pipeline.

#### 6. Optimization Problem Simulation and Response Surfaces

To evaluate optimizer performance under high-dimensional clinical choices, we simulated an 8-site optimization landscape (Problem Size 8, PS 8) with varying site-specific effect sizes. Individual site effects were modeled based on empirical deep brain stimulation (DBS) data [24]assuming that active sites selectively decrease *x*_*base*_ (the desired clinical effect) or yield no effect (Table 1). The corresponding input weighting *b*_*in*_ on *x*_*conflict*_ was fixed to be 10-fold smaller than its effect on *x*_*base*_, consistent with empirical findings that ventral capsule/ventral striatum (VCVS) stimulation predominantly modulates baseline response times. For each problem simulation, the order of the candidate stimulation sites was randomly shuffled. This randomization is critical when evaluating greedy and heuristic optimization algorithms: if the best stimulation site always appears first in the search array, simple greedy strategies will artificially appear highly effective due to positional bias. Randomizing site order across simulation replicates breaks initial ordering dependencies, ensuring that performance metrics reflect true algorithmic search efficiency rather than fortunate contact placement.

**Table 1.** Stimulation effect sizes *b*_*in*_ on *x*_*base*_, in log-seconds) across varying problem sizes.

| Problem size | Stimulation effect |  |  |  |  |  |  |  |
| --- | --- | --- | --- | --- | --- | --- | --- | --- |
|  | 1 | 2 | 3 | 4 | 5 | 6 | 7 | 8 |
| 8 | 0 | -0.005 | -0.01 | -0.02 | -0.03 | -0.031 | -0.04 | -0.07 |

### Performance Metrics

We evaluate optimization performance using two metrics: mean trial-level regret and expected accuracy [28]. In closed-loop DBS tuning, evaluating regret alongside accuracy reveals how algorithms handle observation noise. Because reaction times are highly variable, under-exploratory algorithms risk premature convergence by committing early to suboptimal contacts based on noisy signals. Conversely, exploratory algorithms accept higher initial per-trial regret to average out noise, which is essential for escaping local maxima and identifying the true global optimum.

#### Trial-Level Regret

For a single simulation replicate i, the instant trial-level regret *R*_*i*_ (*t*) at trial t is the penalty incurred by choosing contact *a*_*i,t*_ instead of the true global optimum 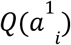 providing information of how far the algorithm was from the optimal.

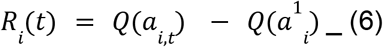

Where 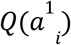 is the ground-truth therapeutic effect of the optimal contact during replicate i, and *Q*(*a*_*i,t*_) is the true effect of the contact selected by the algorithm at trial t.

#### Mean Trial-Level Regret

To evaluate overall search trajectory across varying noise conditions, we average the trial-level regret at each trial t across all N = 1000 independent simulation replicates:

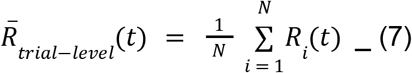

We prioritize mean trial-level regret over cumulative regret because it directly reveals the temporal trajectory and decay of search error, pinpointing the exact trial at which learning saturates. Furthermore, it retains a direct clinical interpretation quantifying the expected reaction-time penalty in seconds at any given trial without obscuring late-stage convergence under an accumulating sum.

#### Recommendation Accuracy

Expected accuracy measures the probability that the algorithm correctly identifies the true optimal contact *a*^1^ at the end of horizon T across all N = 1000 independent runs:

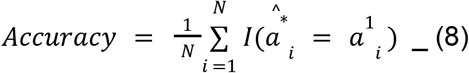

where *I*(.) is an indicator function that equals 1 if the recommended optimal arm 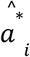 matches the true generator optimal arm 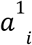, and N = 1000 represents the total number of independent simulation replicates.

### Convergence metrics

Convergence criteria provide a formal framework to evaluate the efficiency and temporal dynamics of the optimization process. By assessing these criteria across trials, we can identify when an algorithm stabilizes and determine whether optimal performance can be achieved with fewer trials, directly minimizing clinical evaluation time and patient burden. Because reaction time (RT) is a noisy biomarker, relying solely on immediate stability risks premature termination driven by transient observation noise. To prevent early convergence, we enforce a mandatory burn-in horizon *T*_*burn*−*in*_ before any dynamic stopping condition can trigger, given by

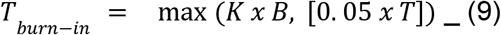

where K is the number of stimulation contacts (arms), B is the block size, and T is the total trial horizon. This burn-in threshold guarantees that every contact is sampled at least once and that initial noise does not cause premature convergence. We evaluate three primary operational stopping criteria:

#### Fixed number of trials

As in [28], a fixed number of trials T is the primary convergence criterion used as a baseline benchmark. The optimization simulation stops once it reaches T.

#### Adaptive Arm Selection Stability

To assess whether the algorithm has reached a behavioral steady state, we track contact allocation over a sliding look-back window W. The window size is scaled to the trial block size (*W* = 10 ∗ *B*). Stability is defined as the empirical probability of selecting the currently favored arm within W.

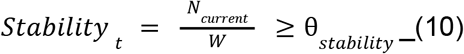

Where *N*_*current*_ represents the selection count within the look back window of the currently active stimulation contact, θ_*stability*_ is the stability threshold (e.g., 0.80 or 0.90). This criterion tests whether the algorithm consistently exploits a single contact target.

#### Relative Empirical Cumulative Reward and Regret Flattening

To determine whether performance has plateaued under variable signal-to-noise conditions, we implement an adaptive thresholding mechanism. Because fixed absolute thresholds fail to account for inter-patient behavioral variability, our threshold scales dynamically with the patient’s localized signal magnitude. We calculate the absolute peak-to-peak range of both the observed Reaction Time and Regret over the last 2W trials. This localized range *d*_*x*_ is compared directly against its adaptive threshold θ_*x*_ over the 2W trials.

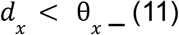

where the localized range is calculated as the peak-to-peak difference over the window:

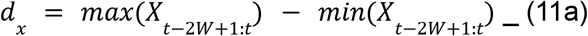

and the adaptive threshold is calculated as:

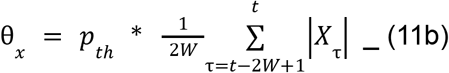

Where *p*_*th*_ is the percentage threshold of the local window’s mean value. The convergence boundary dynamically tightens or widens based on *p*_*th*_.

#### Normalized Global Posterior Precision Variance

For Bayesian algorithms that maintain internal belief distributions over contact effects, we evaluate the system’s informational certainty using a posterior variance criterion. Rather than evaluating performance metrics backward across a temporal look-back window, this mechanism assesses the distribution of parameter uncertainty across all candidate stimulation contacts at trial t. The individual stimulation contact variance are compared directly against to the adaptive variance threshold θ_*v*_ (*t*) to check if the following inequality is satisfied:

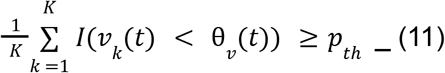

Where K is the total number of stimulation contacts, *v*_*k*_ (*t*)represent the posterior variance of the stimulation contact k at adaptive threshold variance θ_*v*_ (*t*), I(.) is a binary indicator if its true or false, and *p*_*th*_ is the percentage threshold of stimulation contact required to have stabilized (e.g., 0.80). The adaptive variance threshold θ_*v*_ (*t*) is calculated as

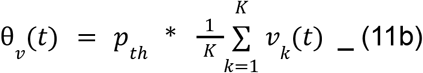

Meeting this criterion flags that a meaningful subset of stimulation contact has reached a high level of certainty relative to the rest of the system.

### Algorithmic Optimization Framework

Algorithmic Optimization Framework

The Multi-Armed Bandit (MAB) optimization framework in this work expands upon our previous formulation [28]. We frame the identification of the optimal Deep Brain Stimulation (DBS) electrode contact on a directional or cylindrical lead as a sequential decision-making minimization problem. Each candidate contact represents a discrete arm *k* ∈ {1, …, *K*}, where the objective is to minimize reaction time (RT) measured during the Multi-Source Interference Task (MSIT).

For baseline comparisons, we retain the foundational mathematical formulations, conjugate updates, and belief dynamics established in our previous work. This baseline suite includes classic heuristic policies greedy and ϵ-greedy strategies as well as standard Thompson Sampling (TS) variants assuming Bernoulli, Poisson, and Normal data likelihoods (denoted as TS-Bernoulli, TS-Poisson, and TS-Normal, respectively), Bayesian Upper Confidence Bound (Bayes-UCB), and Causal Thompson Sampling (C-TS).

To overcome key limitations of standard bandit strategies under non-stationary, noisy clinical observations, we introduce several novel algorithmic formulations and modify existing architectures to improve tracking fidelity and parameter estimation. The mathematical formulations, belief update dynamics, and decision rules for each candidate algorithm are detailed in the following subsections.

#### Upper Confidence Bound (UCB1) and Variants

The standard Upper Confidence Bound (UCB1) algorithm is implemented the same as in [28], but instead of using raw observations, we modified the strategy to normalize rewards using standard global Min-Max normalization. This layer maps all historical reaction times to a standardized [0, 1] metric before computing the selection boundaries to protect the optimization from outlier noise. Further, this ensures proper alignment between the biomarker scale and the exploration bonus, preventing scale mismatches from allowing either term to dominate the optimization process. Furthermore, we present two variations of the UCB: one that provides explicit temporal control over exploration and exploitation, and another that utilizes a variance-dependent version to control it.

#### UCB with Time-Decaying Exploration (UCBTemp)

To explicitly control the transition from exploration to exploitation over time, we introduce a time-decaying variant (UCBTemp). UCBTemp prioritizes exploratory sampling early in the optimization process and systematically reduces exploration width as trials progress. For each candidate contact k, the acquisition cost function evaluated at trial block

To implement control over the exploration-exploitation balance, we developed the UCBTemp. UCBTemp prioritizes exploration early and systematically cools as the trial progresses. The cost function evaluated at each block selection step is defined as

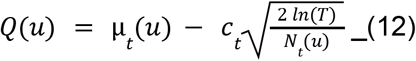

Where μ_*t*_ (*u*) is the vector of the normalized mean reaction time (RT) of stimulation sites in u, *c*_*t*_ is the time decaying exploration rate. The decay is calculated based on the total trial horizon T and the block size B using a configuration parameter ϵ to determine the overall decay exponent *p*.

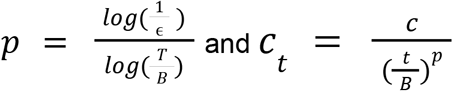

#### Upper Confidence Bound for Variances (UCBTuned)

The UCBTuned overcomes the limitation of treating all arms with identical sample counts equally by dynamically scaling its exploration width based on the empirical variance of each individual stimulation site. This allocates more exploratory trials to contacts exhibiting highly volatile outcomes. The cost function minimized for contact selection is defined as

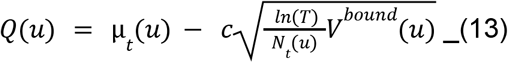

Where α is the configurable tuning parameter, *v*^*bound*^(*u*) is the upper bound for the variance estimate given by

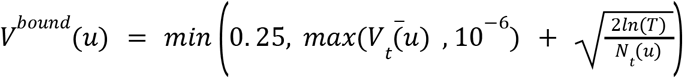

Where 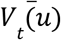 is the normalized stimulation contact variance, lower bounded at 10^−6^ to ensure numerical stability during initial trials. The value 0.25 represents the theoretical maximum variance for normalized [0, 1] rewards, serving as a hard upper bound.

#### Block-Variance Bayes-UCB

At its core, the Block-Variance Bayes-UCB evaluates data at two distinct structural levels. First, it analyzes the internal variance within a block to determine how much individual trials fluctuate around their local mean. Second, it evaluates the block as a single collective unit to measure how far its overall performance has shifted compared to historical data. This approach differs fundamentally from the previous heuristic Bayes-UCB, which relied strictly on a rule-based filter at the macroscopic block level. While this dual-level perspective significantly improves information absorption, it also increases the algorithm’s sensitivity to outlier data, making its exploration boundaries highly susceptible to noisy reaction times (RT).

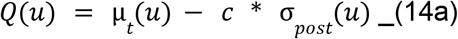

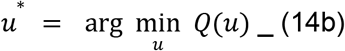

Where u represents a candidate stimulation contact, μ_*t*_ (*u*) is the estimated mean reaction time for that contact, σ_*post*_ (*u*) is the uncertainty boundary, and C is exploration constant. To compute μ_*t*_ (*u*) and σ_*post*_ (*u*), the algorithm tracks both an unknown mean and an unknown variance simultaneously using a continuous Normal-Gamma conjugate prior framework. The posterior update is given by: λ_*t*_ = λ_0_ + *B* where λ is the precision scale, denoting the algorithm’s relative confidence in the estimated mean μ scaled by block size B.

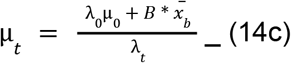

where μ is the estimated mean reaction time of the contact.

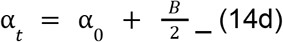

where α is the shape parameter of the posterior distribution.

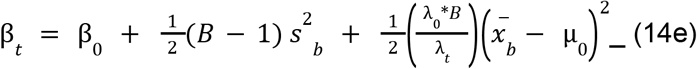

where β is the scale parameter denoting the accumulated variance i.e volatility.

Where sample mean is 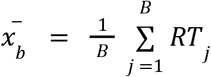 and sample variance is 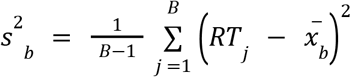

The final analytical uncertainty boundary used in the objective function is computed directly as

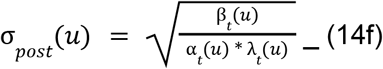

Subscript 0 denotes the prior hyperparameters from past blocks, while Subscript *t* denotes the updated posterior hyperparameters calculated after the current block.

#### Block-Variance Thompson Sampling

The Block-Variance Thompson Sampling framework treats both the expected mean reaction time and the variance as simultaneously unknown, co-dependent variables within a joint Normal-Gamma continuous framework. This differs fundamentally from the TS-Normal algorithm, which assumed an unknown variance but treated the mean reaction time as fixed and known. By removing this constraint, the new implementation allows the algorithm’s belief to adjust to concurrent shifts in both a contact’s Reaction Time (RT) performance and its uncertainty.

It utilizes the same continuous Normal-Gamma conjugate prior as the Block-Variance Bayes-UCB, sharing the same posterior hyperparameter update structure (λ_*t*_, μ_*t*_, α_*t*_, β_*t*_). The core difference lies entirely in its action selection policy. At the beginning of each block, the algorithm draws a random precision(τ) from the posterior Gamma distribution, and subsequently draws a random mean 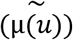 reaction time from a Normal distribution scaled by the exact precision sample.

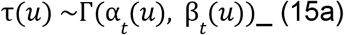

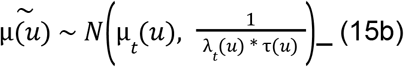

Where τ is sampled environmental precision acting as a temporary measure of the contact’s uncertainty for current selection step, 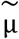 is the final randomized performance metric used to rank and select the next stimulation contact to stimulate at using the selection policy

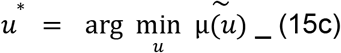

### Algorithm Optimization and Simulation Protocols

To systematically evaluate the optimization frameworks without imposing clinical burdens on human participants, we deployed an *in silico* simulation testbed leveraging the multi-armed bandit (MAB) closed-loop architecture developed in the Generator Model section. This setup is preserved directly from previous work [28] to maintain a consistent benchmarking baseline.

The simulation environment evaluates algorithmic performance across a diverse set of variables. The Signal-to-Noise Ratio (SNR) controls the baseline environmental noise mixed into the reaction time (RT) streams. The Block Size (B) defines the fixed number of trials used for data aggregation and contact selection. The Horizon Trials (T) represent the total number of sequential trials allowed for each optimization run. The ensemble size represents how many times these Horizon Trials (T) are independently repeated to simulate a multi-day timeline. Finally, the Problem Size (PS) establishes the total number of available stimulation contacts in the action space. To achieve rigorous statistical validation and account for random environmental noise, each simulation configuration was run across 1,000 independent replicates.

#### Structural Streamlining: Elimination of the Sensor Model

To evaluate the effects of removal of the state space tracking layer, we simulate data based on the prior sensor based framework [28] where the optimization stimulation contact is suggested and selected based on the latent state. Here, we compare estimated latent cognitive state *x*_*base, sensor*_ with the actual ground truth *x*_*base, generator*_. The core idea of this is to show that while the sensor model is designed to isolate cognitive states, inaccuracies in the model estimation introduces an artificial noise into the control loop, degrading the overall advantage of using the cognitive state.

To test this, we simulated 1,000 independent replicates using a fixed problem size of 8 stimulation sites (PS = 8) and a fixed block size of 15 trials (B = 15). A block size of B = 15 was selected based on the parameter sensitivity analysis, which demonstrated that optimization accuracy plateaus at this threshold, balancing local estimation stability against algorithmic responsiveness. The simulation was configured with a uniformly distributed signal-to-noise ratio (SNR) across realizations to mimic a diverse patient population, matching the validation baseline established in [28]. We performed an ensemble of 1 to 7 days, with each day comprising 600 trials.

Post hoc, we evaluated the structural fidelity between the states, the variance ratio, and overall rank estimation by the optimization algorithms.

##### Structural Fidelity

We calculate the squared Pearson correlation coefficient (*R*^2^) between the *x*_*base, generator*_ and the *x*_*base, sensor*_. This measures how well the sensor tracks the overall shape, peaks, and valleys of the underlying latent cognitive state over time, regardless of whether the absolute values are scaled incorrectly.

##### Variance Ratio

To quantify amplitude distortion due to the use of sensor model, we calculated the unscaled Variance Ratio (*V*_*R*_) between the sensor signal and the generator model. This metric evaluates the preservation of trial-by-trial volatility

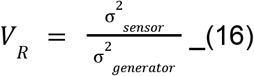

Where 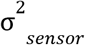 is the variance in the sensor and 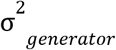 is the variance of the generator. The *V*_*R*_ represents the overall volatility and noise amplification introduced into the data stream by the sensor model.

##### Rank Optimization Failure rate

We calculate average mean values based on *x*_*base, generator*_ and *x*_*base, sensor*_ We then rank the stimulation contacts based on that. This provides a perceived stimulation effect (reward) landscape by the algorithms. We then use sensor values and cross-references it with the true top-performing arm from the ground truth. If they do not match, it is logged as an optimization failure. The final value represents the percentage of total independent runs where the sensor data misled the allocator into selecting a suboptimal arm.

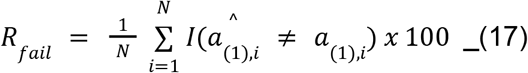

Where I(.) is an indicator function that equals 1 if the sensor-selected optimal arm 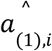 does not match the true generator optimal arm *a*_(1),*i*_, and N = 1000 represents the total number of independent simulation replicates. The final value represents the percentage of total independent runs where the sensor model’s tracking inaccuracies misled the allocation policy into choosing a suboptimal stimulation site.

#### Continuous Optimization Architecture

Prior work in [28]. implemented an ensemble approach of optimization. In this simulation, we implemented a continuous optimization approach to prevent the data resets of the ensemble (day-wise) framework. The baseline 600-trial limit was originally set to model the upper limit of patients performing MSIT trials in a single session. However, our continuous framework evaluates the benefit of saving this data across multiple sessions without resets. The day-wise approach ran separate blocks of T = 600 trials over 1 to 7 days and wiped the history each day. In contrast, the continuous architecture runs uninterrupted across the entire timeline. We set the continuous horizons to 600, 1200, 1800, 2400, 3000, 3600, and 4200 trials. This allowed us to directly match the 1-to-7-day ensemble’s performance metrics. To evaluate this setup, we simulated 1,000 independent replicates for stimulation sites (PS = 8) and a block size of 15 trials (B = 15, balancing local estimation stability and responsiveness). The simulation used a uniformly distributed signal-to-noise ratio (SNR) across the 1,000 realizations. Finally, we compared the performance metrics of both approaches.

#### Convergence Dynamics and Stopping Metrics

We applied the four convergence criteria defined in Section “Convergence metrics” to investigate the possibility of reducing the number of overall trials. Because participants can only tolerate a limited number of trials per session, we tested whether early termination rules could halt the simulation as soon as an algorithm stabilized. To prevent noisy reaction times from causing premature stops, we enforced a burn-in threshold (*T*_*burn*−*in*_ = *PS* × *B*). This guaranteed that no algorithm could stop early until all 8 stimulation sites (PS = 8) were sampled at least once. The block size B was fixed as 15, and the simulation used a uniformly distributed signal-to-noise ratio (SNR) across the 1,000 realizations using raw Reaction Time (RT) as the decision biomarker.

After this burn-in period, a simulation run was halted immediately if it met any of the adaptive criteria. These included arm selection stability, regret flattening, or posterior variance. If no adaptive criteria were met, the run defaulted to the fixed number of trials (T = 4200). Finally, we recorded the total trial count and performance accuracy at termination to analyze the trade-off between potential trial savings and overall performance.

#### Algorithmic Calibration and Hyperparameter Tuning

To find the optimal hyperparameters for each algorithm and ensure a fair comparison evaluation, we evaluated the frameworks using a comprehensive simulation protocol. Every configuration was tested across all seven progressive trial horizons: 600, 1200, 1800, 2400, 3000, 3600, and 4200 trials. All runs utilized raw Reaction Time (RT) as the decision biomarker, maintained a fixed block size of 15 trials (B = 15), and used a problem size of 8 stimulation sites (PS = 8). Finally, each simulation used a uniformly distributed signal-to-noise ratio (SNR) across 1,000 independent realizations.

##### Hyperparameter Grid Search

We used a grid search to find the optimal hyperparameter settings for each algorithm. This calibration ensures that every framework runs at its best. By using these optimized settings, any differences in performance reflect the fundamental mechanics of the algorithms rather than poor parameter choices. These ranges were selected to systematically sweep each algorithm across a spectrum from conservative exploitation to aggressive exploration.

For this grid search, we evaluated each of the five allocation algorithms across 25 discrete, linearly spaced parameter steps (Table 2):

**Table 2.**
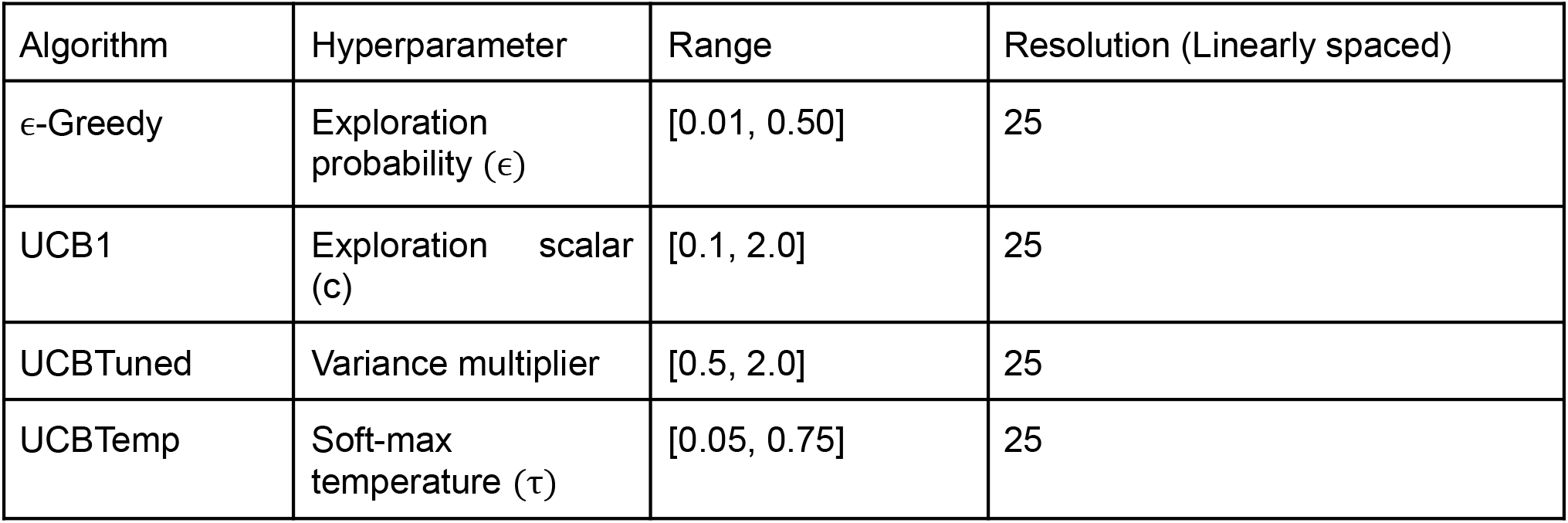
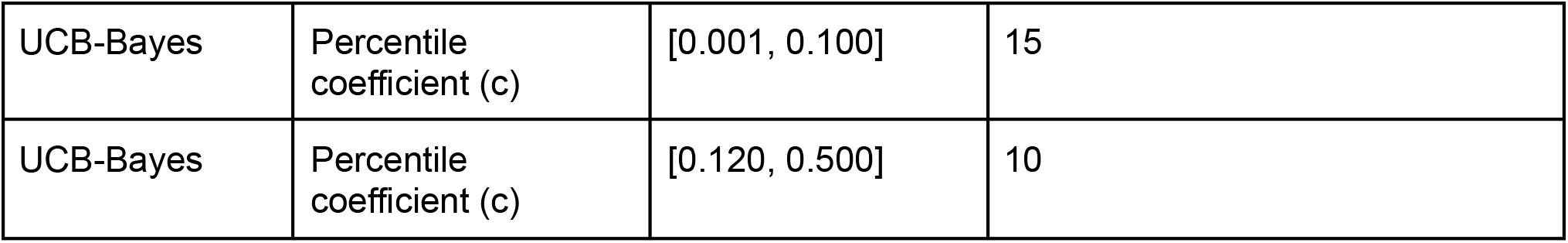
Hyperparameter calibration grid search boundaries and step resolutions. Every configuration was swept over a total of 25 discrete steps (with UCB-Bayes split to sample lower values with higher density) across seven progressive trial horizons (600 to 4,200 trials) using 1,000 independent replicates per configuration.

| Algorithm | Hyperparameter | Range | Resolution (Linearly spaced) |
| --- | --- | --- | --- |
| ε-Greedy | Exploration probability ($\epsilon$) | [0.01, 0.50] | 25 |
| UCB1 | Exploration scalar ($c$) | [0.1, 2.0] | 25 |
| UCBTuned | Variance multiplier | [0.5, 2.0] | 25 |
| UCBTemp | Soft-max temperature ($\tau$) | [0.05, 0.75] | 25 |
| UCB-Bayes | Percentile coefficient (c) | [0.001, 0.100] | 15 |
| UCB-Bayes | Percentile coefficient (c) | [0.120, 0.500] | 10 |

##### Prior Sensitivity Analysis

Alongside the hyperparameter optimization, we conducted a prior sensitivity analysis using the same simulation setup. We wanted to see how our initial baseline assumptions affect the pipeline. Specifically, we tracked whether different starting distributions speed up or slow down how fast the algorithms learn before real data takes over.

To test this, we systematically changed the settings (hyperparameters) of the Normal-Gamma prior distribution. We adjusted the prior reward mean (μ_0_), sample precision (ν_0_), shape (α_0_), and rate (β_0_). Here we compared three specific algorithms: Block-Variance Bayes-UCB, Block-Variance Thompson Sampling, and standard Thompson Sampling (TS-Normal). We systematically tested them across six discrete operational scenarios (Table 3). These specific scenarios were designed to evaluate how the algorithms handle realistic clinical initialization challenges, including miscalibrated expectations (optimistic or pessimistic means) and erroneous confidence levels (excessive or insufficient prior variance).

**Table 3.** Initial parameter configurations and operational conditions for the prior sensitivity analysis. By shifting these starting values across a range of different scenarios, we can isolate exactly how initialization biases affect early-stage exploration and long-term stability.

| Prior scenario | $\mu_0$ | $\lambda_0$ | $\alpha_0$ |
| --- | --- | --- | --- |
| Baseline | 1.5 | 1 | 1 |
| Pessimistic Mean | 0.5 | 1 | 1 |
| Optimistic Mean | 1 | 100 | 1 |
| Overconfident Precision | 1 | 0.01 | 1 |
| High Variance Bias | 1 | 1 | 1 |
| Low Variance Bias | 1 | 1 | 100 |

#### Environmental Stress-Testing

To ensure the closed-loop optimization pipeline can handle real-world clinical volatility, we conducted a set of simulations focusing on environmental stress-testing. In practical deployment, a resilient algorithm must handle two main challenges: enduring extreme signal corruption and adapting to baseline shifts. To measure how well the algorithms recover under this noise and volatility, we tested the multi-armed bandit framework across a maximum horizon of 4,200 trials. Just like our previous tests, these simulations used a fixed block size of 15 trials (B = 15), performed with raw Reaction Time (RT) and a uniformly distributed signal-to-noise ratio (SNR) across 1,000 independent runs. We subjected the framework to two distinct stress configurations:

##### SNR Sensitivity Gradient

We systematically modulated the noise floor of the raw biomarker, the raw Reaction Time (RT), across an extended range from 16dB down to −25dB. Because real-world clinical noise levels are unknown, this gradient helps us identify which specific algorithms perform best across varying noise conditions, and where their internal logic completely breaks down.

##### Non-Stationary Change-Point

To simulate sudden, acute shifts in a patient’s underlying cognitive or physiological baseline, such as medication adjustments or diurnal fluctuations, we introduced an unannounced change-point mid-session. Evaluating this extreme scenario tests the algorithm’s adaptiveness in rapidly unlearning stale reward estimates and re-allocating stimulation, as well as its overall robustness against complete performance breakdown when baseline conditions shift. Although such abrupt baseline inversions represent highly unlikely clinical occurrences during a standard session, evaluating this extreme scenario allows the framework to establish absolute safety boundaries.

#### The Rolling Arena Subspace Tournament Architecture

In practical clinical settings, hardware limitations restrict how many stimulation contacts can be active or configured at the same time. Specifically, implantable neural stimulators are physically constrained by limited active channels on the pulse generator, switching latencies between contacts, and battery power/charge-density safety limits. Because of this constraint, standard multi-armed bandit frameworks suffer from slow convergence speeds as the number of available contacts grows, requiring more trials than a patient can reasonably perform in a single session. This means results from the simulations would not easily translate into real-world practice without a way to scale up. To make our optimization framework useful and scalable under real-world hardware limits, we developed the rolling arena subspace search method.

Instead of optimizing across a large array of contacts all at once, this method restricts the active search space to a smaller, hardware-compatible sub-arena (e.g., four active contacts at a time). The multi-armed bandit algorithm runs normally within this smaller constrained arena, collecting data across a fixed number of trials to evaluate performance. Once the system identifies the worst-performing contact with sufficient statistical confidence, it permanently eliminates it from the active arena. A completely new, untested contact from the remaining pool is then rolled in to take its place. This iterative loop repeats until all available options have been tested. This approach ensures the framework remains highly sample-efficient and well within safe patient trial budgets.

We evaluated these metrics in the fixed-trial execution setup across 12 distinct configurations, testing baseline trial limits ranging from 200 to 1,200 (specifically 200, 300, 400, 450, 500, 600, 700, 800, 900, 1,000, 1,100, and 1,200 trials). Within this setup, initial and final stages are intentionally allocated full trial durations (e.g., 900 trials), while intermediate stages run at half duration (e.g., 450 trials) to ensure sufficient baseline sampling without unnecessary computation. Evaluating these configurations allows us to map the precise trade-off between trial savings and the risk of prematurely losing the best clinical treatment option.

## III. Results

### Elimination of the Sensor Model

Initially, we observed that the tracking error followed the same relative performance hierarchy across behavioral states (Table 4) as established in prior work [28], with slight numerical variations expected due to the stochastic noise across the 1000 replicates. To ensure that the NRMSE bounds were not masking underlying directional mismatches or phase errors, we evaluated the Pearson Correlation Coefficient (*R*^2^,Table 4) as a complementary metric. This mirrored the NRMSE trend demonstrating sensor model ability to track the time varying patterns of the generator.

**Table 4.** Comparative performance metrics across behavioral tracking states (averaged over 1,000 independent replicates). NRMSE and *R*^2^ indicate how closely the sensor model tracks the overall pattern and trajectory of the generator signal. The Variance Ratio *V*_*R*_ indicates whether the sensor model amplifies (*V*_*R*_ > 1, over-responsive) or suppresses (*V*_*R*_ < 1,smoothing) generator variance. The ranking failure rate measures how frequently tracking distortions mislead the optimization algorithm’s decisions.

| State | NRMSE | Pearson correlation ($R^2$) | Variance Ratio ($V_R$) | Ranking Fail Rate (%) |
| --- | --- | --- | --- | --- |
| $RT$ | 0.1069 | 0.8132 | 0.6596 | 11.9% |
| $x_{baseline}$ | 0.0719 | 0.8745 | 1.5174 | 24.5% |
| $x_{conflict}$ | 0.0953 | 0.7585 | 2.4804 | 31.9% |

To investigate the signal’s smoothness and the amplification introduced by the sensor model, we calculated the Variance Ratio (VR). This analysis uncovered a critical tracking issue that was completely missed by relying solely on NRMSE or *R*^2^. Here the sensor’s variance is approximately 51.7% greater than generator’s variance during baseline tracking and 148% greater during conflict state.

This divergence shows that the sensor model amplifies internal high-frequency noise alongside the true generator ground truth. To further understand how this amplified noise influences the overall decision making, we calculated the optimization algorithm’s ranking failure rate. The smoothed, variance-suppressed raw RT signal (*V*_*R*_ =0.6596) maintained the lowest operational disruption, failing in only 11.9% of optimization choices. Conversely, the massive variance inflation found in the latent states severely misleads the selection, culminating in a high ranking failure rate of 24.5% during baseline tracking and 31.9% failure rate under conflict conditions.

We compared the algorithm’s overall accuracy with and without the sensor to show that sensor introduced noise was causing the algorithm to be misled (Figure 2)

**Figure 2.**
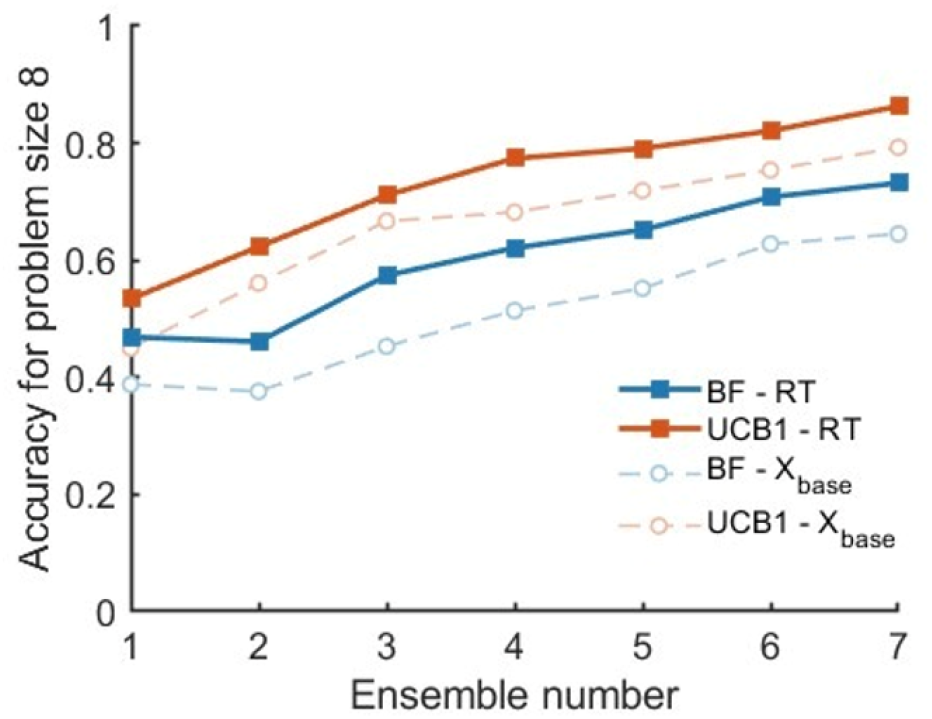
Impact of sensor-model noise on UCB1 optimization performance. A) Algorithmic optimization trajectories over a 4,200-trial timeline (x-axis) comparing performance accuracy (y-axis) for UCB1 operating directly on ground-truth generator data versus sensor-mediated tracking (averaged over 1,000 independent replicates). Evaluating choices directly on clean generator data protects the algorithm from spurious, amplified variance spikes. Bypassing the noisy sensor model eliminates artificial variance inflation, enabling the algorithm to resolve underlying reward trends faster and achieve higher selection accuracy.

**Figure 3.**
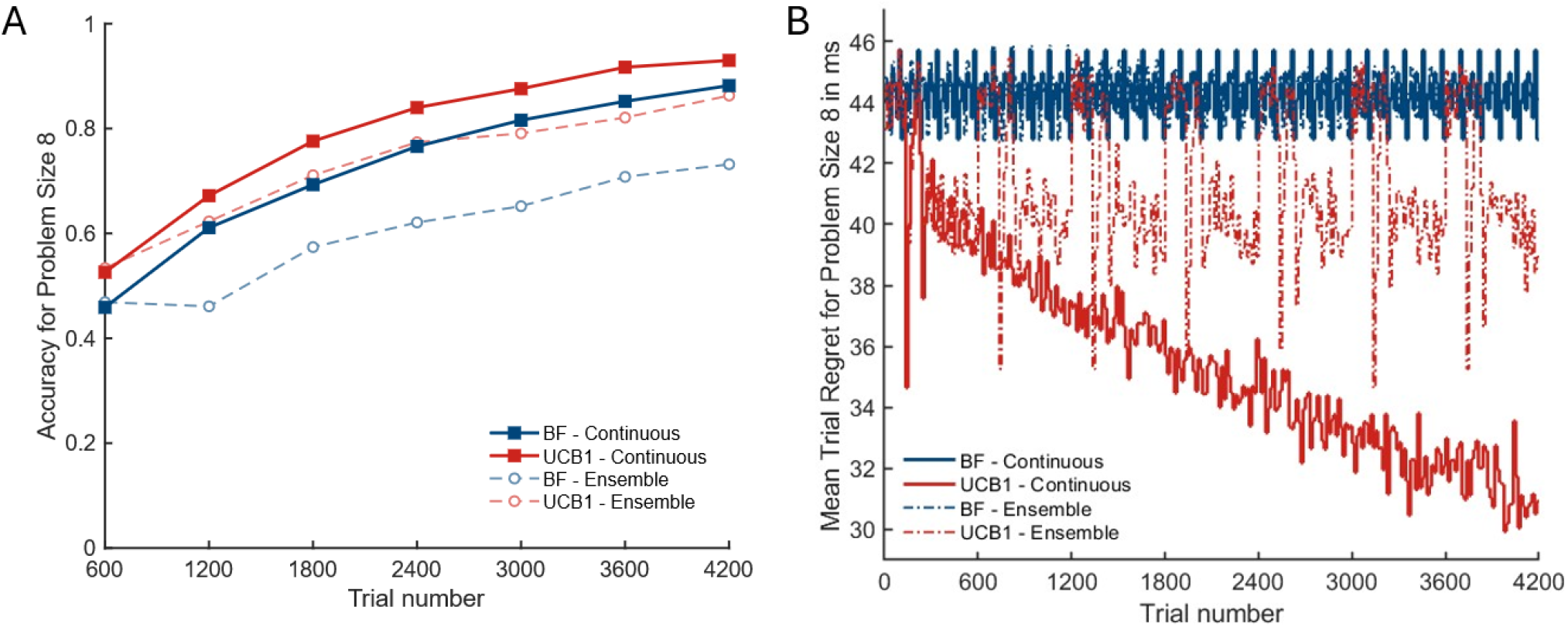
Algorithmic optimization trajectories over a 4200-trial timeline, comparing the continuous optimization process against the resetting day-wise ensemble approach (averaged over 1,000 independent replicates). (A) Selection Accuracy: The continuous framework achieves superior accuracy, whereas the ensemble model plateaus a lower accuracy. (B) Mean Trial-Level Regret: The regret trajectory demonstrates that in the ensemble approach, algorithmic learning is truncated daily (visible from the sudden reset every 600 trials for UCB1-Ensemble) in the ensemble approach, preventing the average mistake rate from dropping below a fixed operational floor. Conversely, the continuous optimization model exhibits a steady downward decay in mean trial-level regret as the number of trials increases, demonstrating a continuous reduction in optimization errors.

### Continuous Optimization Architecture

To investigate whether optimizing over days as a single, uninterrupted process improves algorithmic efficiency, we implemented a continuous optimization framework and evaluated its performance against the ensemble approach [28]. By tracking the system across trials, we observed that consolidating the learning process yields a clear performance advantage. This approach significantly improves both selection accuracy and mean trial-level regret.

The performance divergence stems directly from the number of data samples available to the algorithm. The continuous architecture allows the algorithm to learn from a single growing pool of data. As trials accumulate from 600 up to 4200, the algorithm gains more samples to compute its Reaction Time (RT) estimates. This expanding sample size narrows the confidence intervals around each of the 8 stimulation sites enabling high-confidence decisions. In contrast, the ensemble approach resets the session every 600 trials. This action starves the algorithm of data and truncates its learning process, preventing it from ever accumulating enough samples to minimize its errors.

### Convergence Dynamics and Stopping Metrics

We performed simulations to investigate whether adaptive convergence criteria could reduce the total trial count and reach peak performance faster than a fixed-threshold constraint. We found that the adaptive criteria do not provide an independent learning advantage over the threshold-based approach. As shown in Figure 4, performance accuracy follows an asymptotic trajectory as the number of trials increases. This trend confirms that accuracy depends directly on the sample size and the subsequent information gained by the algorithm.

**Figure 4.**
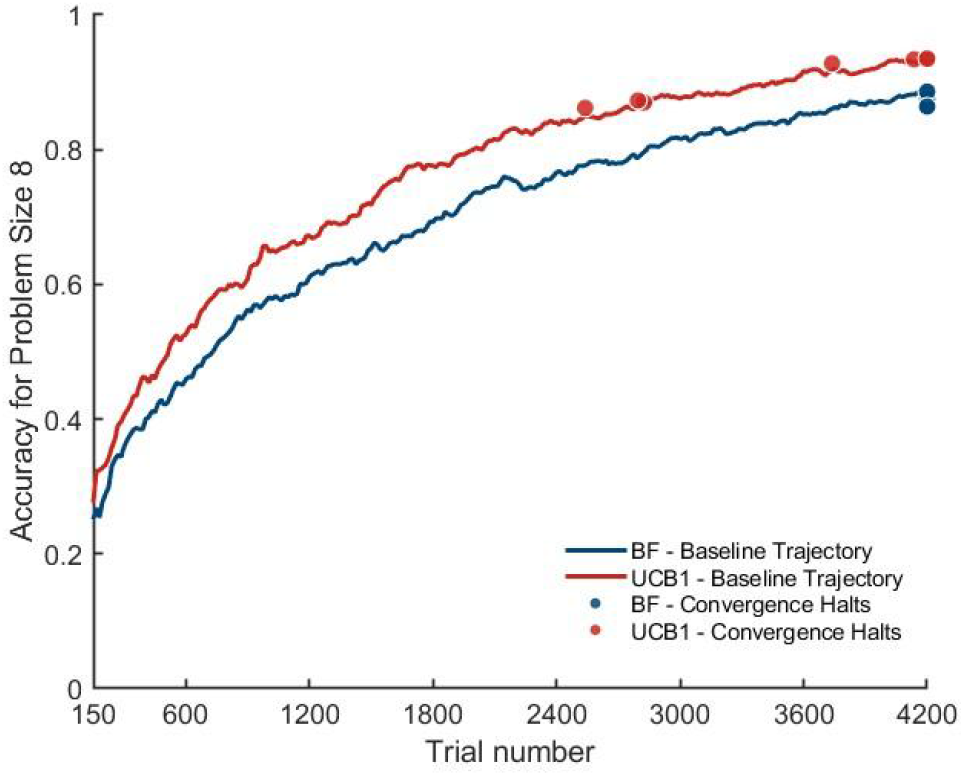
Algorithmic optimization trajectories over a 4,200-trial timeline, comparing adaptive versus fixed-threshold convergence criteria across 8 stimulation contacts (averaged over 1,000 independent replicates). Performance accuracy (y-axis) is plotted as a function of total trials (x-axis) for different convergence criteria, overlaid with the average number of trials (Convergence Halts) required to satisfy each stopping rule. The trajectories demonstrate that adaptive stopping truncates the optimization timeline once localized stability is reached without accelerating the underlying learning rate.

To further evaluate this relationship, we ran simulations that terminated dynamically based on the adaptive stopping rules. These tests confirm that adaptive criteria successfully enable early stopping. However, this early termination introduces a strict trade-off between trial savings and final selection accuracy. The data show that to achieve a high expected accuracy, the algorithm must run for more trials. Adaptive stopping does not accelerate the underlying learning rate; rather, it simply cuts the timeline short once a localized stability threshold is satisfied.

### Algorithmic Calibration and Hyperparameter Tuning

Hyperparameter optimization revealed that tuning the UCB1 exploration constant c improves performance accuracy compared to the standard baseline value (c = 1.0) across problem size 8 (figure 5). The parameter c directly controls the exploration-exploitation balance, where lower values prioritize immediate exploitation and higher values enforce exploration. For c < 0.25, performance accuracy degraded because the algorithm behaved greedily, committing prematurely to initial stimulation contacts. Optimal performance accuracy was achieved within c ∈ [0.30, 0.45], outperforming the standard c = 1.0 baseline previously used as in [28]. Beyond c = 0.45, accuracy progressively decreased as increased exploration weighting caused the algorithm to persistently sample non-optimal contacts despite lower empirical reward evidence.

**Figure 5.**
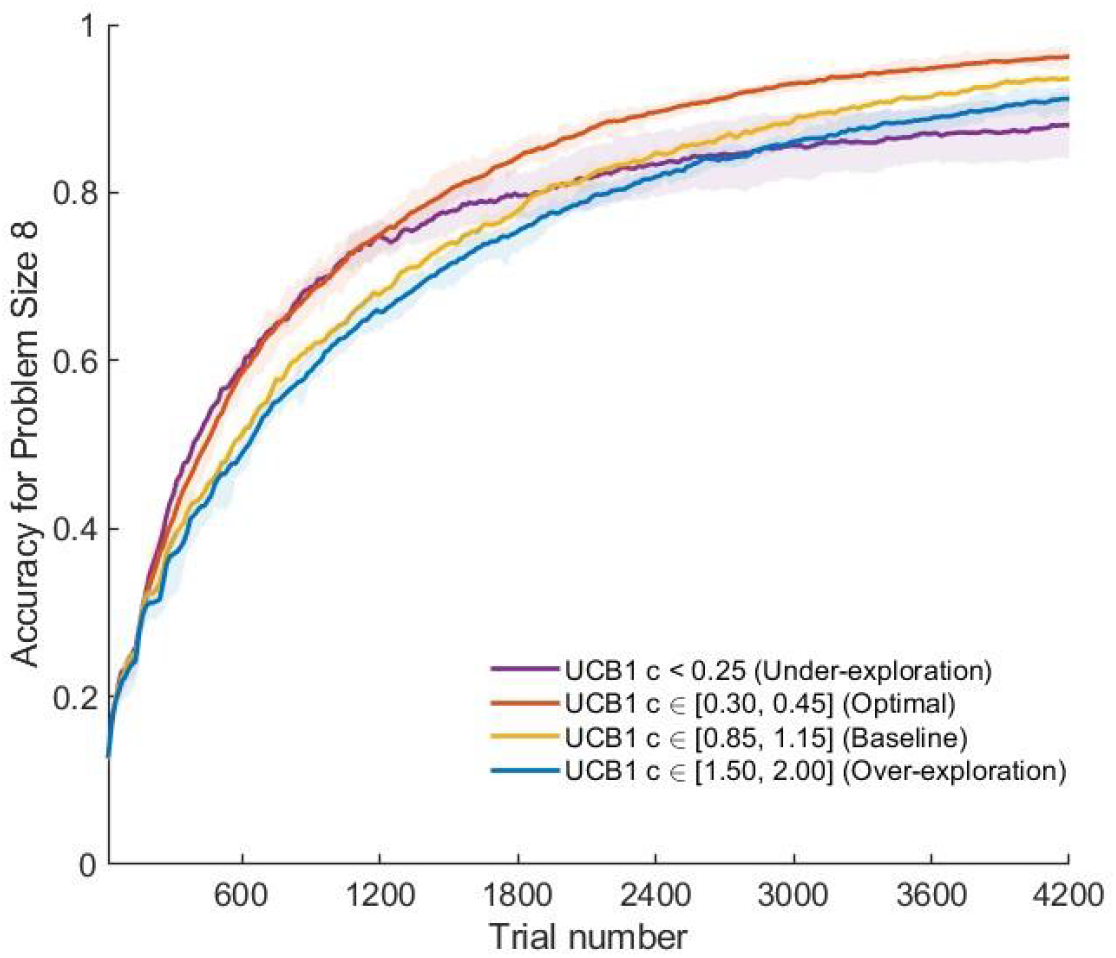
Algorithmic calibration and hyperparameter tuning for the UCB1 exploration constant (c). Algorithmic optimization trajectories over a 4,200-trial timeline (x-axis) for UCB1 under varying exploration constants (c). Trajectories represent mean performance accuracy (y-axis) evaluated across 1,000 independent replicates. Shaded bands denote the range of performance across parameter bins. The optimal hyperparameter range (c ∈ [0.30, 0.45]) achieves higher final accuracy than the standard baseline (c ∈ [0.85, 1.15]), while under-exploration (c < 0.25) and over-exploration (c ∈ [1.50, 2.00]) lead to reduced overall accuracy.

### Prior Sensitivity Analysis

Prior sensitivity analysis revealed that initializing Block-Variance Bayes-UCB with an optimistic mean prior yields superior overall performance accuracy compared to alternative prior configurations (figure 6). Because the optimistic prior assumes high potential across all stimulation contacts (i.e., anticipating a lower reaction time), the algorithm is driven to systematically explore all available options early in the session. Conversely, a pessimistic mean prior assumes contacts are suboptimal, leading to slightly higher exploitation early on. As shown in figure 6, although the pessimistic prior briefly maintains slightly higher performance accuracy initially, its trajectory plateaus around 1,500 trials, after which the optimistic prior permanently surpasses it.

**Figure 6.**
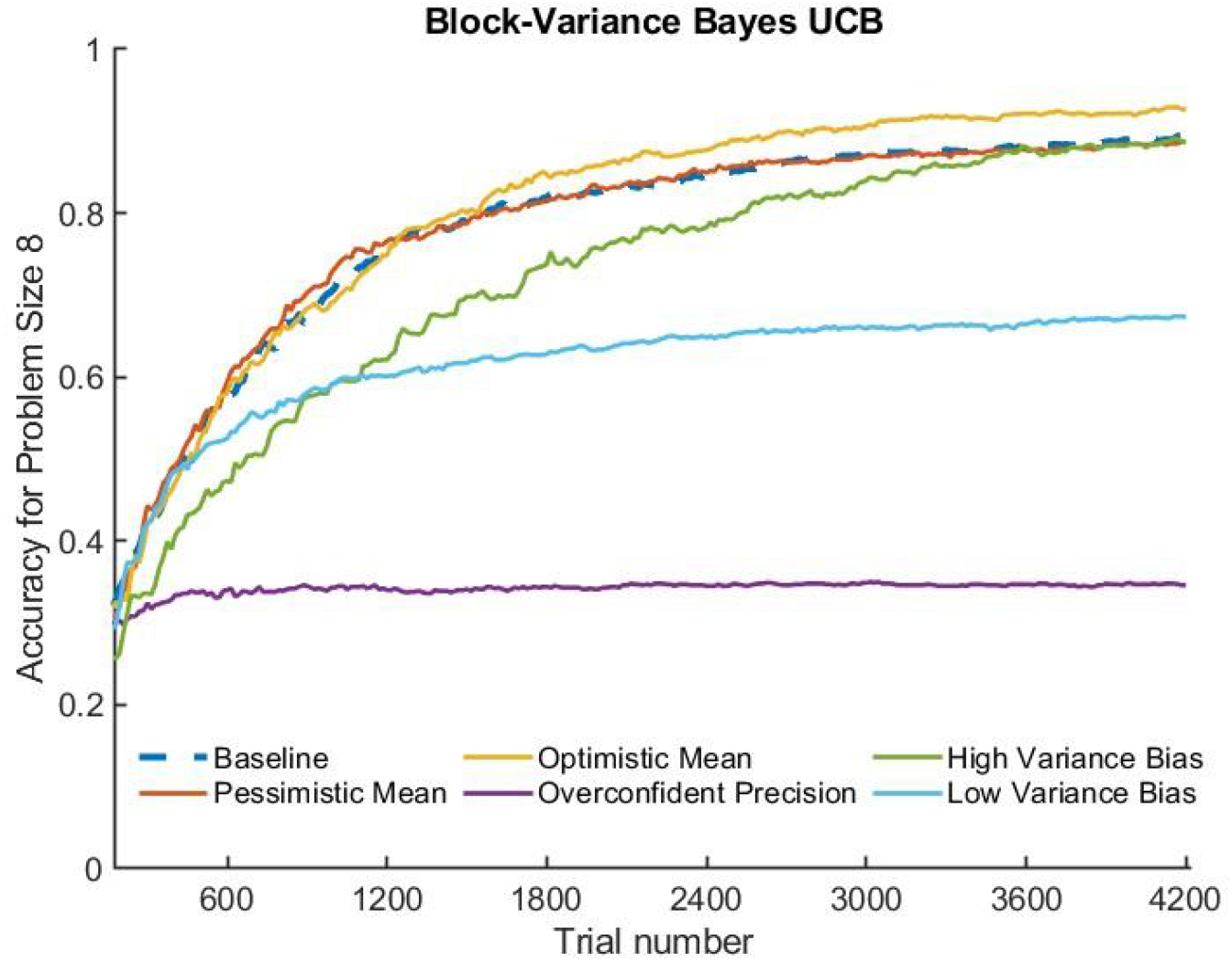
Algorithmic optimization trajectories over a 4200-trial timeline for Block-Variance Bayes-UCB under varying prior assumptions. Trajectories represent mean performance accuracy (y-axis) across total trials (x-axis) evaluated over 1,000 independent replicates. The plot compares baseline, pessimistic mean, optimistic mean, high/low variance bias, and overconfident precision configurations. The optimistic mean prior achieves the highest overall accuracy by promoting systematic early-stage exploration. In contrast, pessimistic, overconfident, and biased variance configurations suffer from premature convergence or persistent over-exploration.

The variance bias configurations further highlight the trade-offs in belief updating. High variance bias assumes an unstable environment, causing the algorithm to continuously mistrust incoming observations and indulge in excessive, persistent exploration; this is reflected in its consistently lower performance accuracy relative to low variance bias. Conversely, low variance bias assumes a stable environment, leading the algorithm to over-trust early samples and exhibit greedy, premature exploitation. Consequently, while accuracy under low variance bias outpaces that of high variance bias early on, performance stalls completely after approximately 800 trials. Finally, the overconfident precision prior places excessive weight on initial assumptions, failing to integrate empirical feedback and exhibiting the lowest overall performance across the timeline. Overall, the optimistic mean prior demonstrates superior long-term optimization accuracy. Notably, baseline priors utilized in [28]. behaved similarly to the pessimistic prior, explaining their lower overall accuracy compared to the optimistic initialization strategy.

### Environmental Stress-Testing

Environmental stress-testing reveals that algorithmic performance accuracy is heavily dependent on the signal-to-noise ratio (SNR) of the system (Figure 7). In low-SNR conditions where environmental volatility and measurement noise are high the performance accuracy of Brute Force declines significantly due to its non-adaptive, unguided sampling scheme. Conversely, UCB1 consistently outperforms Brute Force across both high-SNR and low-SNR regimes. By dynamically balancing exploration and exploitation, UCB1 mitigates the corrupting effects of observational noise. These results demonstrate that adaptive learning algorithms provide a far more robust and reliable framework for identifying optimal stimulation contacts than standard exhaustive search strategies, particularly in noisy clinical environments.

**Figure 7.**
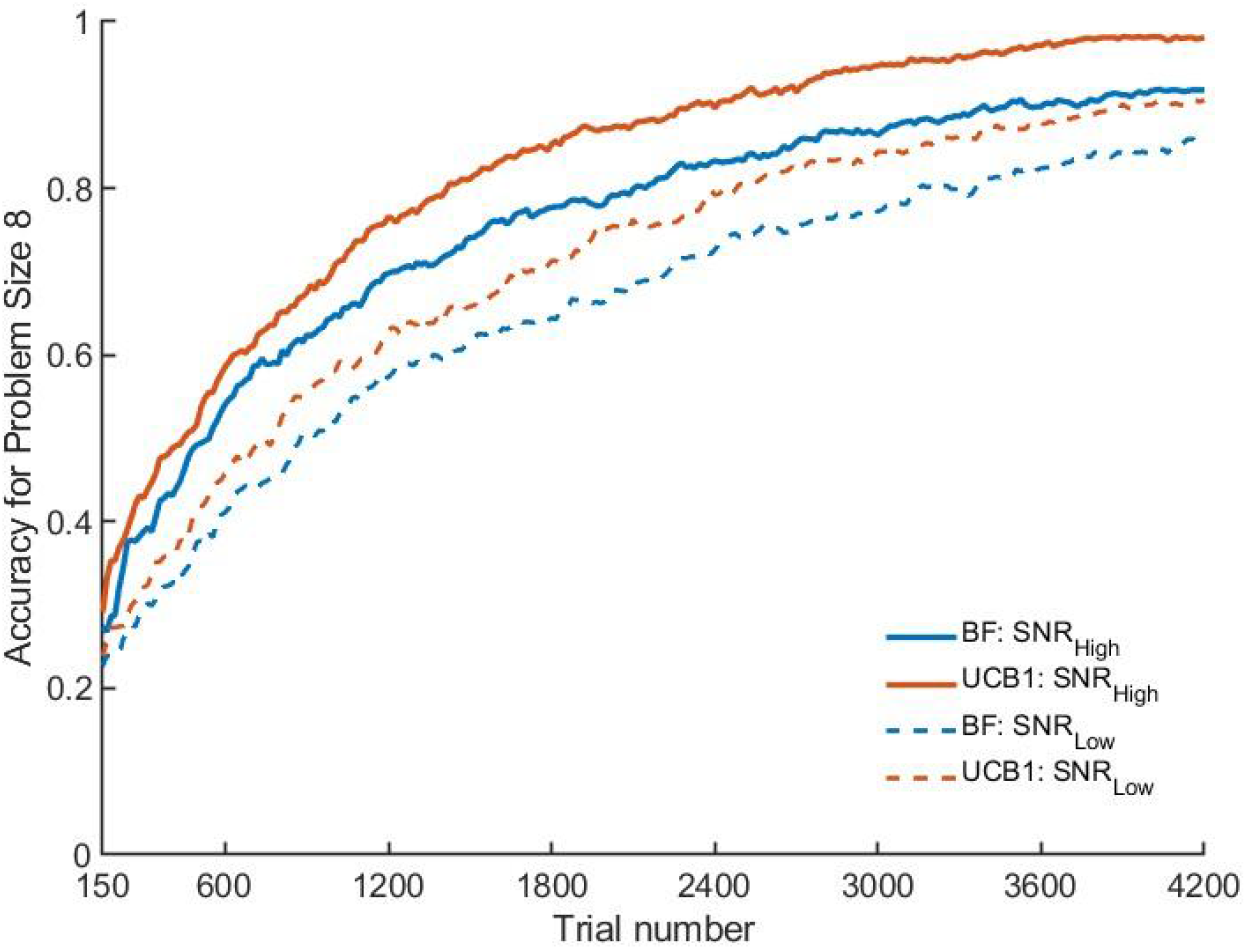
Environmental stress-testing under varying signal-to-noise ratios (SNR). Algorithmic optimization trajectories over a 4200-trial timeline (x-axis) comparing Brute Force (BF, solid lines) and UCB1 (dotted lines) across high- and low-SNR conditions. Trajectories represent mean performance accuracy (y-axis) evaluated across 1000 independent replicates. In low-SNR regimes, UCB1 maintains robust selection accuracy, whereas non-adaptive Brute Force degrades significantly under observational noise.

### Non-Stationary Change-Point

To evaluate algorithmic robustness under biological non-stationarity, we introduced an unannounced, acute change-point at Trial 2,100, where the underlying performance of the optimal and sub-optimal stimulation contacts was completely inverted (Figure 8). Prior to the change-point, all algorithms tend to the initial optimal contact, maintaining high baseline accuracy. Immediately following the shift, performance accuracy experiences a sharp drop as the historical policy is rendered obsolete.

**Figure 8.**
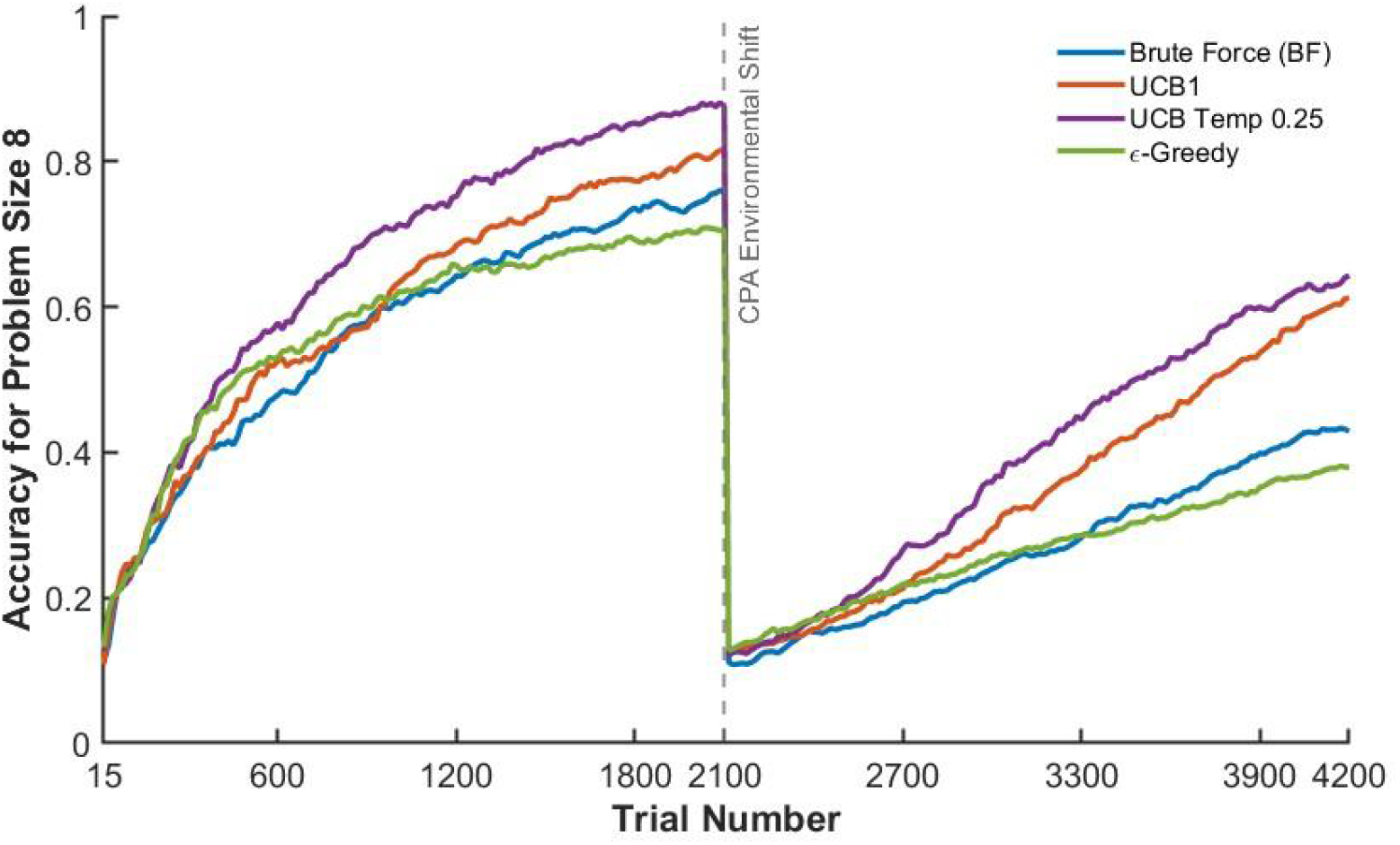
Algorithmic adaptation under a non-stationary change-point. Optimization trajectories over a 4,200-trial session (x-axis) comparing performance accuracy (y-axis) for Brute Force, UCB1, UCB Temp 0.25 and ϵ-greedy across 1000 independent replicates. An unannounced, acute baseline inversion is introduced mid-session (Trial 2100) to simulate a sudden shift in cognitive baseline (e.g., medication onset or fatigue). Trajectories illustrate each algorithm’s ability to detect structural shifts, reset exploration boundaries, and re-identify the new optimal stimulation contact.

Rather than remaining permanently locked into the former top-performing site, all algorithms rapidly detect the shift, causing a sharp spike in uncertainty that triggers a new phase of dynamic re-exploration. As shown in Figure 8, both UCB1 and UCB Temp 0.25 quickly recover from the inversion, systematically identifying the new optimal contact and re-establishing high performance accuracy. This stress-test confirms two vital properties: first, it establishes a clear safety boundary, proving that the algorithms can recover from extreme baseline disruptions; second, it confirms that the framework tracks genuine structural physiological changes rather than simply overfitting to transient, trial-to-trial noise.

### The Rolling Arena Subspace Tournament Architecture

To address the high-dimensional search space problem, which often arises from hardware limits on how many contacts can be active at once, or from the dynamic re-introduction of contacts previously deemed unusable, we evaluated algorithmic performance within a Rolling Arena Subspace Tournament framework. By dynamically cycling contacts between the active “arena” and the benched pool, the framework restricts the active evaluation fold without losing track of potential candidates.

As illustrated in Figure 9, optimization within the rolling arena achieves overall performance accuracy comparable to a continuous, full-space optimization across all eight contacts simultaneously. Interestingly, post-2,000 trials, a slight performance degradation is observed for UCB Temp 0.25 and ϵ-greedy, likely due to their fixed exploration parameters struggling with late-stage arena rotation. In contrast, Brute Force and standard UCB1 exhibit a subtle performance improvement over the same period as lower-performing contacts are systematically benched. Ultimately, the subspace tournament framework demonstrates that high-dimensional contact arrays can be efficiently optimized without sacrificing long-term accuracy, stability, or clinical robustness.

**Figure 9.**
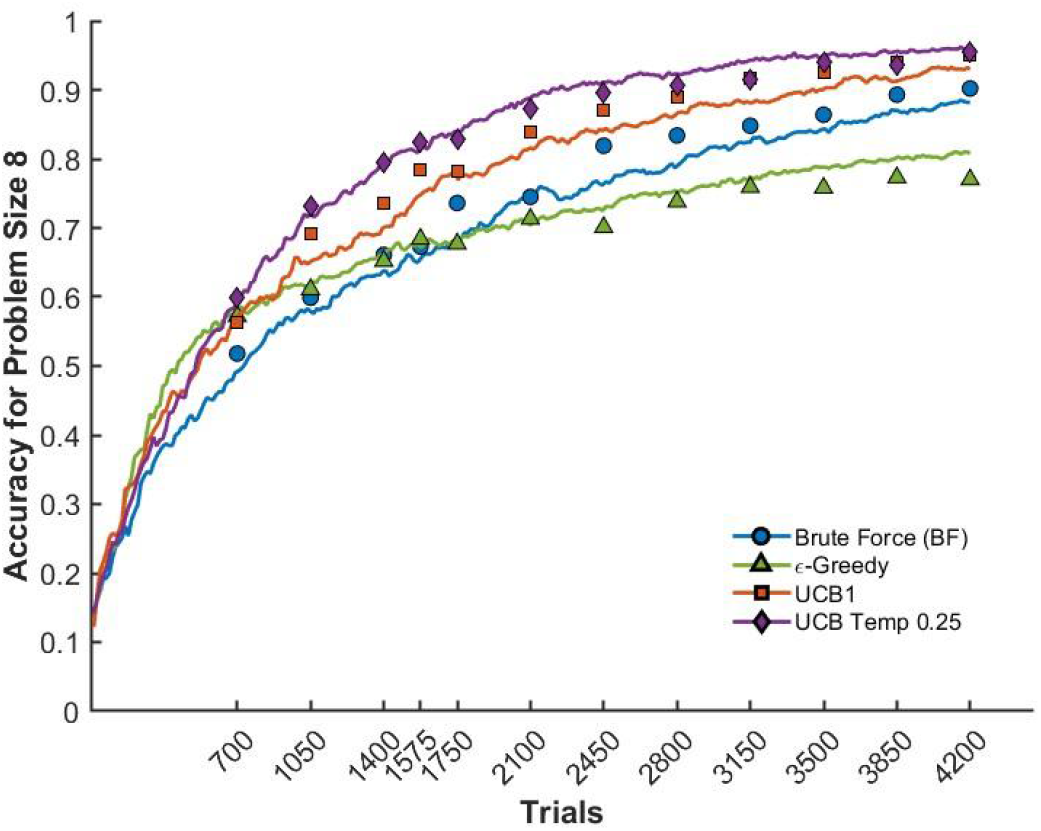
Algorithmic optimization under a Rolling Arena Subspace Tournament framework. Optimization trajectories over a 4,200-trial session (x-axis) comparing performance accuracy (y-axis) for Brute Force, UCB1, UCB Temp 0.25 and ϵ-greedy across 1,000 independent replicates. Solid lines depict baseline performance accuracy across a continuous optimization over the full problem size of 8 contacts, while distinct marker shapes represent the performance accuracy observed within the rolling arena framework. The framework dynamically manages a large pool of candidate stimulation contacts by evaluating a restricted, active subset (the arena) while demoting underperforming contacts to the benched pool. Trajectories demonstrate rapid early-stage convergence and sustained accuracy comparable to unconstrained, full-space search models.

## IV. Discussion

This study evaluated the algorithmic resilience, convergence dynamics, and clinical viability of different algorithms under noisy and non-stationary conditions. The primary finding demonstrated that the use of a simplified closed loop optimization approach with an optimization of stimulation contacts performed over a continuous trials yields better performance accuracy and regret as compared to the prior closed loop approach with intermediary sensor model and discrete discontinuous optimization [28], we show that this is caused by the use of noisy estimation of the sensor model.

### Observational Noise and Sensor Model Elimination

A primary insight from our investigation is that intermediate sensor models can inadvertently corrupt optimization decisions if they amplify internal variance. While conventional metrics like NRMSE and Pearson’s *R*^2^ suggested the sensor model tracked the generator signal well, the Variance Ratio (*V*_*R*_) revealed a severe hidden vulnerability. Furthermore, our post hoc analysis reveals that, in latent baseline and conflict states, the sensor model inflated variance by approximately 51.7% and 148% relative to the ground truth generator signal, respectively. This high-frequency noise amplification severely misleads the optimization algorithm, resulting in ranking failure rates as high as 31.9%.

By eliminating the intermediate sensor model and allowing algorithms like UCB1 [29] to interface directly with raw behavioral data (i.e., reaction time), we completely removed these artificial noise artifacts.To be clear, an idealized, highly accurate sensor model would theoretically enhance optimization performance accuracy by providing a denoised representation of underlying cognitive states. However, given the limitations of current estimation methods, present-day sensor models yield noisy approximations that degrade system performance. Bypassing these intermediate modeling layers eliminates “unknown-unknown” noise corruptions, providing a streamlined, highly reliable approach that is significantly more practical for near-term clinical deployment.

### Information Accumulation: Continuous Architecture vs. Daily Resets

In [28], we previously implemented a discrete, discontinuous ensemble approach that reset the optimization process at the start of each session, under the assumption that routine resets were necessary to accommodate potential day-to-day shifts. However, our current comparative simulations show that this daily resetting strategy imposes a severe learning penalty by repeatedly truncating the algorithm’s accumulated sample pool and trapping mean trial-level regret above a fixed operational floor.

By consolidating learning into a single continuous framework, the algorithm is provided with a steadily growing pool of data across trials (expanding from 600 up to 4,200 samples). This sustained information accumulation allows the algorithm more time to learn, progressively narrowing decision confidence intervals around candidate contacts to achieve higher selection accuracy. Crucially, our Change-Point Analysis (CPA) demonstrates that hard resets are not required to handle non-stationary shifts; even if an optimal contact changes mid-stream, continuous adaptive algorithms naturally detect the resulting surge in uncertainty and dynamically re-converge without needing to throw away historical data.

### Convergence Dynamics and Adaptive Stopping

Determining the optimal session endpoint is critical in clinical practice to avoid under-sampling noisy behavioral data while preventing unnecessary patient burden and fatigue. Our findings demonstrate that adaptive and fixed-threshold stopping rules yield equivalent performance accuracy when evaluated over an equal number of trials. While adaptive convergence metrics are useful for early halting when time is strictly limited, truncating a session inherently reduces expected performance accuracy across all algorithms tested. Given that the primary objective is to identify the most effective stimulation setting, utilizing the complete observation window remains preferable whenever clinically feasible.

Here, we identified hyperparameters and priors to ensure the algorithms are optimized before clinical deployment, ensuring each algorithm operates at its expected peak potential rather than under default values. Because committing to a full observation window demands patient engagement, every trial within a session must be maximally informative. Our calibration highlights the critical necessity of balancing exploration and exploitation in clinical optimization tasks.

If an algorithm is configured too greedily, it prematurely locks onto the first moderately effective stimulation contact, abandoning the search before discovering the true global optimum. Conversely, excessive exploration causes the model to degenerate into an unguided brute-force search, repeatedly testing sub-optimal contacts without ever stabilizing on a solution. Establishing calibrated parameters and optimistic priors ensures that observed performance disparities stem from genuine differences in algorithmic capability rather than sub-optimal tuning. This provides a robust foundation for evaluating how these algorithms withstand real-world clinical noise and underlying physiological shifts.

Further clinical viability of these algorithms was evaluated by stressing the environment and isolating their performance across varying Signal-to-Noise Ratios (SNRs). Remarkably, even under high-noise conditions, adaptive bandit algorithms consistently demonstrated superior performance compared to unguided brute-force search. While brute-force methods spend excessive trials repeatedly sampling noisy, sub-optimal contacts, adaptive algorithms effectively filter behavioral noise to converge on the true target. This proves that algorithmic guidance remains robust and highly advantageous even when signal quality degrades.

In clinical practice, patient reaction times are inherently noisy, and the therapeutic efficacy of stimulation contacts can fluctuate over time due to impedance changes, electrode migration, or shifting fatigue levels. To evaluate clinical feasibility under these dynamic conditions, we simulated an extreme scenario where all underlying contact effects were abruptly shuffled midway through the trial sequence. Although such a complete redistribution is unlikely in routine clinical workflows, it serves as a rigorous worst-case stress test for algorithmic adaptability. The adaptive algorithms successfully detected and tracked these abrupt shifts, pivoting to identify the new optimal contact despite the change in underlying response dynamics. This shows that all algorithms are resilient to the environment shifts.

While the prior experiments demonstrated algorithmic resilience, the practicality and scalability of these findings were further confirmed by the Rolling Arena (RA) paradigm. The Rolling Arena framework addressed a core practical challenge in clinical practice: the inability to optimize all stimulation contacts simultaneously within a single go. By evaluating candidate contacts in constrained subsets over time, this paradigm proved that optimization across an eight-contact array remains clinically viable without demanding full, concurrent access. Furthermore, the Rolling Arena demonstrated the key advantage of modular flexibility, allowing new stimulation contacts to be dynamically integrated into the optimization fold as treatment progresses. Together, these experimental evaluations confirm that the adaptive algorithms not only maintain robustness against unpredictable physiological shifts, but also scale seamlessly to real-world clinical constraints. Together, these results demonstrate that adaptive learning algorithms provide a safe, scalable, and clinically viable solution for real-time stimulation contact optimization.

## V. Study Limitations

While these in-silico experiments demonstrate clear algorithmic proof-of-concept, several key clinical translation considerations warrant discussion. Moving toward full clinical deployment involves expanding the multi-dimensional parameter space (amplitude, pulse width, and frequency), incorporating explicit safety boundaries, integrating refined sensor modeling, and exploiting spatial correlations across electrode contacts.

First, the proposed framework focuses solely on selecting the optimal discrete stimulation site. In full clinical practice, DBS optimization requires tuning a broader, multi-dimensional parameter space that includes continuous variables like amplitude, pulse width, and frequency [30]. However, attempting to optimize all continuous and discrete variables simultaneously causes a combinatorial explosion, leading to impractically long convergence times. Our focus on contact selection enables a two-stage solution to this challenge. Because contact location primarily determines target engagement, our framework can first anchor the optimal site. Continuous parameters can then be fine-tuned in a separate, secondary step, simplifying the search space by decoupling the discrete dimension.

Second, the current optimization framework relies exclusively on reaction time as its sole objective metric, operating without explicit safety bounds or penalties for adverse side effects. While reaction time provides a reliable objective measure, optimizing a single behavioral proxy limits real-world clinical utility. In targets like the ventral capsule/ventral striatum (VCVS), stimulation can trigger acute affective side effects, including anxiety, volatility, or mania. Without explicit safety constraints, the optimizer risks selecting settings that improve task performance at the expense of patient safety or psychological comfort. Clinical deployment will therefore require multi-objective reward structures, along with the ability to dynamically prune adverse-effect-producing options from the search space. Transitioning to a weighted optimization framework will allow algorithms to explicitly penalize negative outcomes while incorporating patient-reported preferences, such as mood ratings, as secondary decision factors.

Third, this implementation omitted the construction or re-estimation of an explicit task-specific sensor model. Avoiding custom sensor estimation simplifies the pipeline and reduces computational burden. However, relying on a simplified observational mapping limits state estimation accuracy. If an accurate sensor model is available, or if more advanced modeling techniques are applied, incorporating this mapping could enhance latent state estimation. Implementing a well-calibrated sensor model represents a powerful pathway to improve overall optimization accuracy and convergence speed in future iterations.

Fourth, the framework treats stimulation contacts as independent options, ignoring spatial correlation across electrode sites. In physical DBS leads, neighboring contacts share overlapping electric fields and engage contiguous neural tissue. By treating each contact independently, the algorithm discards valuable structural information. Incorporating spatial kernels or Gaussian Process priors to model performance across lead geometry would exploit this spatial continuity. This approach could significantly reduce search space complexity and accelerate convergence to the optimal contact.

## VI. Conclusion

In summary, we have demonstrated a robust, direct-learning proof-of-concept framework for closed-loop DBS contact optimization using reaction time during cognitive control as a measure of effective target engagement. By eliminating noisy intermediate sensor models and transitioning to a continuous learning architecture, this framework achieves superior selection accuracy and stability under low signal-to-noise ratios, non-stationary physiological shifts, and high-dimensional rolling arena constraints. Future work will focus on expanding optimization across multi-dimensional continuous parameter spaces (amplitude, pulse width, and frequency) while incorporating multi-objective safety boundaries to manage stimulation-induced adverse events. Ultimately, translating these adaptive algorithms into routine psychiatric care will require robust clinical software integration and rigorous usability testing, representing a critical next step toward fully autonomous, patient-tailored DBS programming.

## Acknowledgments

Research reported in this publication was supported by the University of Minnesota’s MnDRIVE (Minnesota’s Discovery, Research and Innovation Economy) initiative, the Minnesota Medical Discovery Team on Addiction, the University of Minnesota Interdisciplinary Doctoral Fellowship, the Dr. Ralph and Marian Falk Medical Research Trust, and the National Institutes of Health / NIH Blueprint (R01MH123634, R01MH119384).

## Conflict of interest

ASW is a consultant for Abbott and has unlicensed intellectual property related to deep brain stimulation. ASW holds equity in Resilient Neurotherapeutics. SSN report no biomedical financial interests or potential conflicts of interest.

## Data and Code Availability Statement

The extracted data from simulations and code that support the findings of this study are openly available at the following URL/DOI: https://github.com/tne-lab/dbs-accuracy-optimization-2026

## Notes

https://github.com/tne-lab/dbs-accuracy-optimization-2026

### Competing Interest Statement

The authors have declared no competing interest.

